# Contrasting signatures of introgression in North American box turtle (*Terrapene* spp.) contact zones

**DOI:** 10.1101/752196

**Authors:** Bradley T. Martin, Marlis R. Douglas, Tyler K. Chafin, John S. Placyk, Roger D. Birkhead, Christopher A. Phillips, Michael E. Douglas

## Abstract

Hybridization occurs differentially across the genome in a balancing act between selection and migration. With the unprecedented resolution of contemporary sequencing technologies, selection and migration can now be effectively quantified such that researchers can identify genetic elements involved in introgression. Furthermore, genomic patterns can now be associated with ecologically relevant phenotypes, given availability of annotated reference genomes. We do so in North American box turtles (*Terrapene*) by deciphering how selection affects hybrid zones at the interface of species boundaries and identifying genetic regions potentially under selection that may relate to thermal adaptations. Such genes may impact physiological pathways involved in temperature-dependent sex determination, immune system functioning, and hypoxia tolerance. We contrasted these patterns across inter- and intra-specific hybrid zones that differ temporally and biogeographically. We demonstrate hybridization is broadly apparent in *Terrapene*, but with observed genomic cline patterns corresponding to species boundaries at loci potentially associated with thermal adaptation. These loci display signatures of directional introgression within intra-specific boundaries, despite a genome-wide selective trend against intergrades. In contrast, outlier loci for inter-specific comparisons exhibited evidence of being under selection against hybrids. Importantly, adaptations coinciding with species boundaries in *Terrapene* overlap with climatic boundaries and highlight the vulnerability of these terrestrial ectotherms to anthropogenic pressures.

## 1. INTRODUCTION

Hybrid zones are natural laboratories that allow the genetic architecture of local adaptation and/or reproductive isolation to be examined. They frequently juxtapose with underlying ecological gradients, allowing researchers to quantify the how selection impacts the genome (Barton & Hewitt 1985; Payseur 2010). Here, selection might prevent introgression at loci underpinning crucial adaptations while the rest of the genome essentially homogenizes (Via 2009; Feder *et al.* 2013). Once such loci are identified, the phenotypes inferred by adaptive divergence can then be inferred via a bottom-up, “reverse-ecology” approach (Li *et al.* 2008; Tiffin & Ross-Ibarra 2014). Hybrid zones effectively become unique “windows” into the speciation process by allowing functional loci to be associated with various aspects of ecology (Taylor *et al.* 2015).

Diminishing costs associated with genomic sequencing, coupled with an upsurge in genomic annotations, has facilitated the reverse-ecology approach. Examples include adaptive divergence in seasonal growth and variability in immune responses (Rödin□Mörch *et al.* 2019), proactive responses to environmental gradients (Keller & Seehausen 2012; Guo *et al.* 2016; Waterhouse *et al.* 2018; Teske *et al.* 2019), and an upsurge in contemporary effects such as anthropogenic modulation of reproductive boundaries (Garroway *et al.* 2010; Taylor *et al.* 2014; Grabenstein & Taylor 2018). Importantly, researchers can now effectively gauge the how the genome is impacted by ecological and climatic shifts, and the how species distributions are promoted by pre-existing adaptive gradients (Rosenzweig *et al.* 2008; Taylor *et al.* 2015; Ryan *et al.* 2018). Thus, genome-scale datasets can often refine our perspectives on two major evolutionary patterns: First, reproductive boundaries among historically co-existing species can become blurred due to substantial environmental change, a situation directly analogous to the imposition of contact among otherwise allopatric taxa (Rhymer & Simberloff 1996). Second, selection and migration are balanced within hybrid zones (Key 1968; Parmesan *et al.* 1999), and this balance can shift due to rapid and/or exceptional climate change (Seehausen *et al.* 2008; Kearns *et al.* 2018). Genomic data extracted from hybrid zones may thus allow species boundaries to be defined according to their phenotypic and genetic underpinnings.

Species boundaries can either be strengthened (Ryan *et al.* 2018) or eroded (Muhlfeld *et al.* 2014) due to temperature shifts, a major component of climate change, potentially shifting species distributions and/or hybrid zone dynamics. Furthermore the effects of temperature on physiological and cellular mechanisms are well known (Kingsolver 2009), directly affecting growth, development, reproduction, locomotion, and immune response (Keller & Seehausen 2012). As a result, the manner by which thermal gradients interact with species boundaries has become a major focus (Qin *et al.* 2013). Herein, we attempt to clarify how species boundaries reflect environmental processes by quantifying the geographic and ecological foundations of two hybrid zones in the ectothermic North American box turtles (*Terrapene*).

### 1.1 Hybridization in North American box turtles

North American box turtles (Emydidae, *Terrapene*) are long-lived, omnivorous, and primarily terrestrial ectotherms, with a rectangular appearance defined by a dome-shaped dorsal carapace and a ventral plastron hinged to tightly close against the carapace (hence the common name) (Dodd 2001). Their North American range is characterized by two well-known zones of hybridization (Milstead 1969; Dodd 2001; Cureton *et al.* 2011) that provide excellent models from which to contrast regional patterns of hybridization and introgression. To do so, we evaluated four southeastern taxa [the Woodland (*T. carolina carolina*), Gulf Coast (*T. c. major*), Three-toed (*T. carolina triunguis*), and Florida (*T. bauri*) box turtles (Auffenberg 1958, 1959; Milstead & Tinkle 1967; Milstead 1969; Martin *et al.* 2013; Iverson *et al.* 2017)], and two midwestern (the Ornate box turtle, *T. ornata ornata* and *T. c. carolina*; Cureton *et al.* 2011). Each hybridizes regionally, and therefore we focus on inter- and intra-specific contacts within these two regions.

One focal hybrid zone is nested within southeastern North America (Ricketts 1999), where box turtles inhabit a biodiversity hotspot. Here, clear-cutting, invasive species, and altered fire regimes are widespread (Stapanian *et al.* 1997, 1998; van Lear & Harlow 2002), and impact numerous endemic species (Lydeard & Mayden 1995). The region also displays clinal intergradation (i.e., interbreeding between subspecies), as well as hybridization across a variety of taxa (Remington 1968; Swenson & Howard 2004), due largely to coincident ecological and climatic transitions (Swenson & Howard 2005).

By contrast, contact zones in midwestern North America seemingly stem from secondary contact (i.e., resumption of interbreeding following a geographic separation), as associated with postglacial recolonization/ expansion (Swenson & Howard 2005). Here, prairie-grassland habitat has been anthropogenically fragmented such that niche overlap now occurs between grassland and woodland species (Johnson 1994; Samson & Knopf 1994; Rhymer & Simberloff 1996; Samson *et al.* 2004). Furthermore, while overlapping forms in the Midwest represent distinct species, southeastern forms are taxonomically in flux. Species status has varied for *T. m. triunguis*, and *T. c. major* is now viewed as an intergrade population (Butler *et al.* 2011; Iverson *et al.* 2017), despite recent molecular work suggesting specific status for *T. m. triunguis* and phylogenetic structure in *T. c. major* (Martin *et al.* 2013, 2014, 2020). For the sake of clarity, we follow Martin *et al.*, regarding *T. m. triunguis* as a full species and *T. c. major* as a recognized subspecies.

We used ddRAD sequencing (Peterson *et al.* 2012) to contrast genome-wide patterns of clinal introgression within each hybrid zone. We then identified/ quantified loci potentially under selection by mapping them to an available genomic reference and accordingly interpreted their potential ecological associations, which provide invaluable insights into how *Terrapene* has responded to a fluctuating climate. As such, our results extend to a proactive management paradigm that underscores the conservation of co-occurring forms.

## 2. MATERIALS AND METHODS

### 2.1. Tissue and DNA collection

Tissues for *T. carolina*, *T. ornata*, and *T. mexicana triunguis* were collected by volunteers and agency collaborators (Table S1). Additional samples were provided by numerous museums and organizations. Live animals were sampled non-invasively (e.g., blood, toenails, or toe-clips), whereas road-kills were sampled indiscriminately. Isolation of genomic DNA was performed using DNeasy Blood and Tissue Kits (QIAGEN), QIAamp Fast DNA Tissue Kit (QIAGEN), and E.Z.N.A. Tissue DNA Kits (Omega Bio-tek). The presence of genomic DNA was confirmed via gel electrophoresis using a 2% agarose gel.

### 2.2. Library preparation

*In silico* digests were carried out to optimize restriction enzyme selection, using available genomic references [Painted turtle (*Chrysemys picta*), GenBank Accession #: GCA_000241765.2 (Shaffer *et al.* 2013); Fragmatic (Chafin *et al.* 2018); genome size=2.59 × 10^9^ bp]. The distribution of fragments (from N=24 samples) were first optimized then evaluated using an Agilent 4200 TapeStation. Library preparation was conducted per standard protocol (Peterson *et al.* 2012), using *PstI* (5’-CTGCAG-3’) and *MspI* (5’-CCGG-3’) restriction enzymes. We digested ~500-1,000ng of DNA/ sample at 37°C, with unique DNA barcode and sequencing adapters subsequently ligated. Prior to sequencing, quality control checks were performed at the core facility, to include fragment analysis for confirmation of correct size range and quantitative real-time PCR. Individuals (N=96) were pooled per lane of single-end Illumina sequencing at the University of Oregon Genomics and Cell Characterization Core Facility (Hi-Seq 4000, 1×100bp). Populations were randomized across multiple lanes to mitigate batch effects.

### 2.3. Assembly and quality control

Read quality was quantified using FastQC v. 0.11.5, then demultiplexed and aligned using ipyrad v. 0.7.28 (Eaton & Overcast 2020), with reads mapped to the scaffold-level *T. mexicana triunguis* reference assembly (GenBank Accession #: GCA_002925995.2) at a distance threshold of 0.15. Non-mapping reads were discarded. This alignment is herein termed the “scaffold alignment” to differentiate it from a separate transcriptome-mapped alignment (see section 2.6). Barcodes and adapters were trimmed, as were the last five base pair (bp) of each read. Those exceeding five bases with low PHRED quality score (<33) were discarded, and potential paralogs were filtered by excluding loci with high heterozygosity (>75%) or >2 alleles per individual. Loci with a sequencing depth of <20X per individual or <50% presence across individuals were also discarded. Our mapping and filtering steps above were conducted in Ipyrad.

### 2.4. Assessing Admixture and Population Structure

Admixture (Alexander *et al.* 2009) was used to assess contemporary hybridization. It employs a model-based ML approach that estimates the proportion of ancestry shared across the genome-wide average of each sample. *K=*1-13 were used for datasets containing all sequenced taxa and subsets from the Southeast and Midwest hybrid zones, with 20 independent replicates per *K* (AdmixPipe; Mussmann *et al.* 2020). Hierarchical partitioning was done because Admixture often underestimates *K* by detecting only the uppermost hierarchy of population structure (Evanno *et al.* 2005). SNP data were pre-filtered using VCFtools (Danecek *et al.* 2011), with SNPs randomly thinned to one per locus to alleviate linkage bias and filtered by removing sites with a minor allele frequency (MAF)<1.0% to reduce bias associated with erroneous genotypes and singletons (Linck & Battey 2019). Model support across *K*-values was assessed using five-fold cross-validation (Alexander *et al.* 2009). AdmixPipe output was summarized using the Clumpak server (Kopelman *et al.* 2015), with each individual subsequently plotted as a stacked bar-chart (Rosenberg 2004).

We also performed Discriminate Analysis of Principal Components (DAPC) using the *adegenet* R-package (v2.0-0) with identical filtering parameters (Jombart *et al.* 2010). The *find.clusters()* function was utilized with 1,000,000 iterations to determine the optimal *K* with the lowest Bayesian Information Criterion (BIC). DAPC cross-validation (100 replicates, 90% training dataset) then evaluated which principle components and discriminant functions to retain, with individuals plotted against the top three DAPC axes.

Finally, we also ran TESS3 (tess3r R package; Caye *et al.* 2016) to estimate ancestry coefficients (as with Admixture), but also incorporate spatial proximity into ancestral genotype estimates. The TESS3 input alignments were subsets of those used in Admixture to include only individuals with GPS coordinates. Cross-validation (with 10% sites randomly masked) was performed for K=1-10 with twenty independent runs to assess optimal *K*. The output Q-matrix was interpolated using spatial kriging (Jay *et al.* 2012).

### 2.5. Identifying Hybrids

NewHybrids (Anderson & Thompson 2002) was used to assign statistically-supported hybrids to genotype frequency classes (i.e., Pure, F_1_, F_2_, and backcrosses between F_1_ and parental types). The *getTopLoc()* function in HybridDetective (Wringe *et al.* 2017a) reduced the data to 300 loci containing the highest among-population differentiation (*F*_ST_) and lowest linkage disequilibrium correlation (*r*^2^<0.2). Burn-in was 500,000 MCMC generations followed by 2,000,000 post burn-in sweeps. Seeds were randomized and the analysis employed the Jeffrey’s prior for θ and π. To train the data, individuals sampled outside the focal hybrid zones with Admixture proportions=100% were pre-assigned as parentals. The following combinations of taxa were employed: *Terrapene carolina carolina* X *T. c. major*, *T. c. carolina* X *T. m. triunguis*, *T. c. major* X *T. m. triunguis*, and *T. c. carolina* X *T. o. ornata*. A posterior probability threshold >0.8 was required for assignment into the genotype frequency classes, as determined using a power analysis conducted with HybridDetective and ParallelNewHybrid pipelines (Wringe *et al.* 2017b; a).

### 2.6. Genomic Clines among Scaffolds and mRNA Mapping

In addition to the “scaffold alignment,” ipyrad was rerun with reads mapped to the *T. m. triunguis* reference transcriptome (GenBank Accession: GCA_002925995.2), with identical filtering parameters. Three subsets of the resulting “transcriptomic alignment” were generated to retain only individuals per each pairwise combination of southeastern taxa (*T. c. carolina*, *T. c. major*, *T. m. triunguis*). The “scaffold” and “transcriptome” alignments were then independently examined for patterns of differential introgression using Introgress (Gompert & Buerkle 2010) and Bayesian Genomic Clines (Gompert & Buerkle 2012). Both generate genomic clines, which assess locus-specific ancestry to identify outliers *versus* the genome-wide average and can identify outlier SNPs having cline shapes divergent from neutral expectations. Parental reference populations were determined *a priori* via population structure and NewHybrids results. Parental status was considered only for samples with Admixture ancestry coefficients=100%.

Introgress derived neutral expectations from 1,000 parametric simulations (Gompert & Buerkle 2010). Genomic clines were only generated for SNPs with a high allele frequency differential (δ) between parental types (Andrés *et al.* 2013). Outlier SNPs were defined using a Bonferroni-corrected α-significance threshold.

Prior to running BGC, sites with a MAF<5% were removed because an over-abundance of uninformative loci inhibited parameter convergence. Five independent BGC runs were conducted for both scaffold and transcriptomic alignments, which included 1,000,000 and 1,800,000 burn-in, respectively, each with 200,000 post-burn-in generations. Samples were thinned with every 50 iterations retained to mitigate auto-correlation. Genotype uncertainty corrections were applied to each locus, with the sequencing error rate prior computed in ipyrad (ranging from 0.1-0.2%). The BGC linkage model was tested but found to be computationally intractable (i.e., >1Tb memory and unreasonable running times). Upon run completion, parameter traces were visually inspected for convergence. Replicate runs were subsequently combined.

Two parameters [i.e., genomic cline center (α) and rate (ß)] represented BGC output (Gompert & Buerkle 2011). The α-parameter indicates the direction of introgression, with negative and positive outliers depicting excess P_1_ and P_2_ ancestry, respectively. ß characterizes the rate of change, with negative values reflecting a wider genomic cline where loci more freely introgress, and a steeper cline indicated by positive values with relatively sharp transitions from P_1_ to P_2_ ancestry. BGC outliers were considered significant if they met either of two criteria: 1) the 95% credible intervals for α or ß did not overlap zero, or 2) the median of the posterior distribution exceeded the probability distribution’s quantile interval [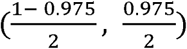 (Gompert & Buerkle 2011)].

### 2.7 Mapping BGC Outliers to Chromosomes

BGC parameters α and ß were plotted onto assembled chromosomes to visualize the distribution of outliers across the genome. The *Terrapene* reference is a scaffold-level assembly, so Minimap2 (Li 2018) and PAFScaff (https://github.com/slimsuite/pafscaff) were used to map the *T. m. triunguis* reference to a closely related chromosome-level assembly (*Trachemys scripta*; Simison *et al.* 2020; GenBank accession: GCA_013100865.1). The ASM20 minimap2 preset was chosen to accommodate the ~5-10% sequence divergence expected between *Trachemys* and *Terrapene* (Feldman & Parham 2002). The mapping connected *Terrapene* BGC outliers with *Trachemys* chromosomal positions, allowing the putative BGC outlier locations to be visualized at the chromosome-level, with plots subsequently generated (Rideogram R package; Hao *et al.* 2020).

### 2.8 Correlating Outliers with Environmental Variables

We also independently identified SNP outliers by environmental association using redundancy analysis (RDA). Outlier SNPs were correlated with the standard WorldClim v2 Bioclimatic variables that included 19 raster layers of temperature and precipitation, plus mean annual solar radiation, mean annual wind speed, and elevation (Fick & Hijmans 2017). The finest available scale (30 arc-seconds) was chosen for each raster. Layers were projected to WGS84 and cropped to the sampling extent (raster R package; Hijmans & Van Etten 2016). Raster values at each sampling location were then extracted.

Each predictor variable was scaled, centered, and tested for normality with a Shapiro-Wilks test. Non-normal distributions were transformed using the bestNormalize R package (Peterson & Cavanaugh 2019) per RDA’s assumptions. The environmental layers were assayed for predictive capabilities with uncorrelated variables retained (adespatial R package, forward selection with 10,000 permutations). Predictors that failed forward selection were removed. To account for underlying spatial influence, distance-based Moran’s eigenvector maps (dbMEM) were generated using sample coordinates (*quickMEM()* R function; Borcard *et al.* 2018). The dbMEMs are a matrix of axes that capture spatial patterns from multiple angles rather than just a latitudinal or longitudinal vector. To reduce overfitting, informative, non-redundant dbMEM axes were subset, using forward selection with 1,000 permutations. Finally, the SNP matrix was imported into R using adegenet, and missing data were imputed as the most frequent allele per population, following RDA assumptions.

A partial RDA (pRDA) conditioned on the dbMEM spatial matrix was then conducted using genotypes as the response variables, with 1,000 permutations (vegan R package; Oksanen *et al.* 2019). This approach “partialed out” spatial autocorrelative effects that could yield false negative SNP-environment associations. Significant RDA axes were determined using vegan’s *anova.cca*() function, and SNPs with loadings +/− 3 standard deviations from the mean on a significant axis were considered outliers. A full RDA and a pRDA conditioned on environment were also conducted to estimate the contributions of spatial *versus* environmental predictors. Each SNP was then correlated pairwise with all environmental variables using Pearson’s correlation coefficient (*r*), and those with the strongest correlations represented the best supported SNP-environment association.

## 3. RESULTS

A total of 368 individuals (Tables S1, S2) were retained across 12,052 (combined alignment), 10,338 (Midwest-only), and 11,308 (Southeast-only) unlinked reference-mapped loci. This, a result of quality control steps and post-alignment filters that eliminated individuals with >90% missing data, and sites with MAF<1.0%. The scaffold alignment included 134,607 variable and 90,777 parsimoniously informative sites. The transcriptome-guided alignment contained 2,741 biallelic SNPs across 247 individuals, with subsets generated for *T. c. carolina* X *T. c. major* (EAxGU), *T. c. carolina* X *T. m. triunguis* (EAxTT), and *T. c. major* X *T. m. triunguis* (GUxTT).

### 3.1. Admixture across the hybrid zones

The combined east/ west Admixture CV (Fig. S1) supported *K*=6 (x□ = 0.19279, SD = 0.00017), followed by *K*=4 (x□ = 0.19490, SD = 0.0016) and *K*=5 (x□ = 0.19738, SD = 0.00013). The analysis indicated population structure for *T. c. carolina*, *T. m. triunguis*, two distinct *T. c. major* subpopulations (Alabama/ Mississippi and Florida panhandles), and northern and southern *T. o. ornata* subpopulations from Illinois+Wisconsin+Iowa and Kansas+Texas+Colorado+Nebraska (Fig. S2). *Terrapene bauri* was excluded due to limited sampling (N=4). Admixture occurred between *T. c. carolina* X *T. c. major* (EAxGU), *T. c. carolina* X *T. m. triunguis* (EAxTT), *T. c. major* X *T. m. triunguis* (GUxTT), *T. o. ornata* X *T. c. carolina* (EAxON), and the two *T. o. ornata* subpopulations.

The lowest CV score for the southeastern Admixture (N=259 individuals) was at *K*=4 (x□ = 0.21851, SD=0.00016), with K=3 (x□ =0.22134, SD=0.00015) and K=5 (x□ = 0.22519, SD=0.00082) trailing (Fig. S3). Southeastern taxa included *T. c. carolina*, *T. c. major*, and *T. m. triunguis*, and their analysis concurred with the all-taxa dataset in terms of both population structure and admixed taxa (Figs. 1, S4). Admixture primarily occurred throughout Alabama and the Florida panhandle (EAxGU), Georgia and South Carolina (EAxTT), and Mississippi/ southern Alabama (GUxTT). The two *T. c. major* subpopulations in the all-taxa analysis were corroborated. The same four southeastern groups plus *T. bauri* were also produced by DAPC (*K*=5; Fig. S5). We found *T. bauri* highly differentiated along axis 1 (71.9% variance explained), whereas axes 2-3 delineated the remaining southeastern taxa (17.5% and 5.73%, respectively).

**Figure 1:**
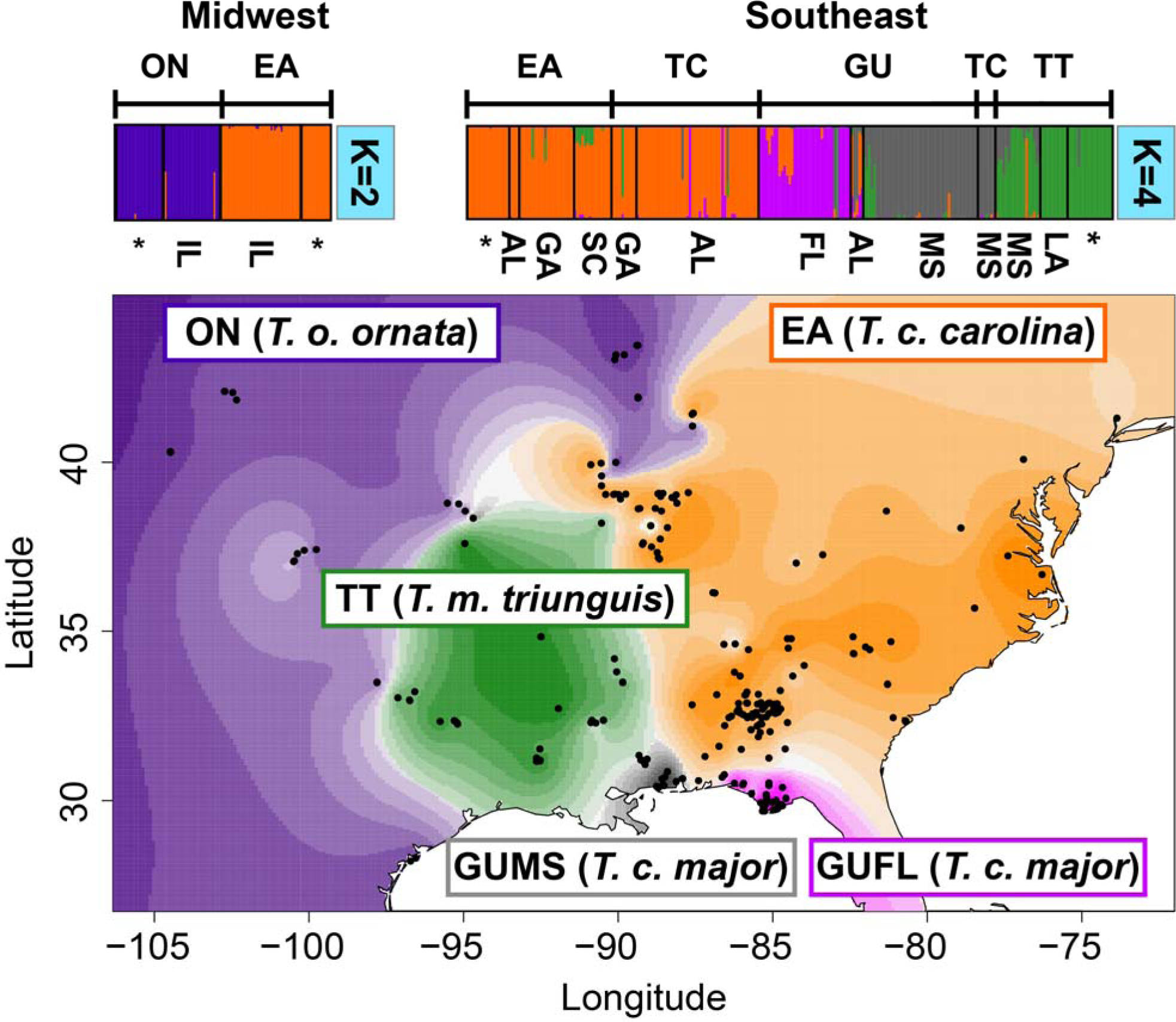
Admixture (barplots) and TESS3 (map) analyses for 11,308 *Terrapene* ddRADseq loci. All plots include individuals (N=320; black circles on map) from the Midwest and Southeast hybrid zones, with the optimal number of clusters (*K*) determined via cross-validation across 20 independent runs. Labels above Admixture plots indicate subspecies (when available) as identified in the field: EA=Woodland (*T. carolina carolina*), ON=Ornate (*T. o. ornata*), TC=*T. carolina* (subspecies identification unavailable), GU=Gulf Coast (*T. c. major*), TT=Three-toed (*T. mexicana triunguis*). Bottom labels show hybrid zone localities by U.S. state: IL=Illinois, AL=Alabama, GA=Georgia, FL=Florida, MS=Mississippi, LA=Louisiana. A * represents a group of “pure” individuals from multiple localities outside the hybrid zones. TESS3 ancestry coefficients (Q) are predicted across the spatial surface via Kriging interpolation (θ=10) and are color-coded with the Admixture plots. Lighter/ darker gradient shades depict lower/ higher Q.

The midwestern analysis (Figs. 1, S6) included an optimal K=2 (x□ = 0.24069, SD= 0.00023), followed by K=3 (x□ =0.25703, SD=0.00454) and K=4 (x□ =0.25861, SD=0.00374) (Fig. S7). The *K*=2 groups consisted of *T. c. carolina* and *T. o. ornata*, with only a few individuals indicating Admixture. At *K*=3, *T. c. carolina* from Illinois split as a distinct group, although only a few of the Admixture proportions approached 100%. At *K*=4, the northern and southern *T. o. ornata* subpopulations produced by the all-taxa analysis also materialized.

TESS3 corroborated both midwestern and southeastern Admixture analyses, with *T. o. ornata*, *T. c. carolina T. m. triunguis*, and the two *T. c. major* subpopulations in Alabama/ Mississippi and Florida being delineated (Fig. 1). The Kriging interpolation also spatially highlighted ancestry gradients consistent with Admixture (FIG. S8), with lower surface prediction scores concordant with areas that contained frequently mixed ancestry.

### 3.2. Genealogical hybrid classification

HybridDetective confirmed convergence for inter- and intra-simulation replicates (Fig. S9) with 500,000 burn-in and 2,000,000 post-burn-in sweeps (the EAxON analysis required 4,000,000 sweeps with 1,000,000 burn-in). Our power analyses suggested 90% assignment accuracy (+/-SD) for all genotype classes at a critical threshold of 0.8 (Figs. S10, S12, S14, S16). Statistical power was also elevated (≥0.8), although some genotype classes for EAxGU displayed relatively lower power (<0.8) (Figs. S11, S13, S15, S17).

The southeastern hybrid zone consisted entirely of backcrosses (F_1_ hybrids X parental types), F_2_-generation, and unassigned (>F_2_) hybrids (Fig. 2, Table S3). Specifically, the EAxGU analysis identified backcrosses with parental *T. c. major* and F_2_ hybrids in the Florida panhandle and southern Alabama (Fig. 2A). Similarly, all hybrid-generation EAxTT individuals from Georgia were identified as backcrosses with both parental types, whereas South Carolina hybrids were backcrosses with *T. c. carolina* (Fig. 2B).

**Figure 2:**
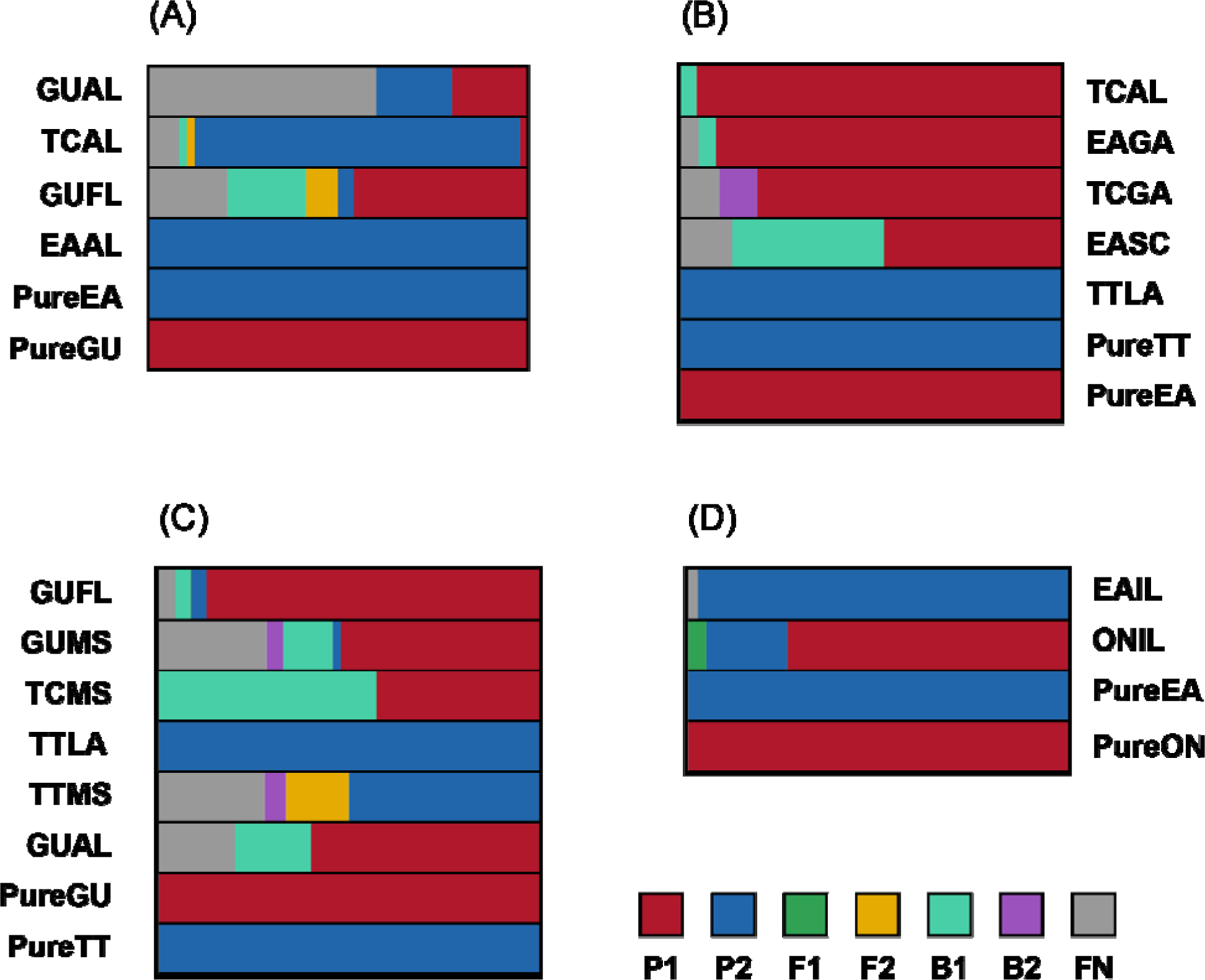
Population-level NewHybrids plots for four pairs of southeastern and midwestern *Terrapene* taxa. Individuals were collapsed into populations based on field identification at the subspecific level. The first two characters represent: GU=Gulf Coast, *T. c. major*; EA=Woodland, *T. c. carolina*; TT=Three-toed, *T. m. triunguis*; ON=Ornate, *T. o. ornata*; TC=*T. carolina* (subspecies-level field identification unavailable). The last two characters represent U.S. state: AL=Alabama, FL=Florida, MS=Mississippi, SC=South Carolina, GA=Georgia, LA=Louisiana, IL=Illinois). Each plot corresponds to tests between parental groups (A) EAxGU (N=109), (B) EAxTT (N=135), (C) GUxTT (N=139), and (D) EAxON (N=112). A posterior probability threshold >0.8 was required for genotype frequency class assignments, which included P_1_ and P_2_ (parental types), F_1_ and F_2_ (first and second-generation hybrids), backcrosses (B_1_ and B_2_), and F_N_ (unclassified).

Second-generation hybrids and backcrosses with both parental types were evident among GUxTT (Fig. 2C). Mississippi contained individuals with all three hybrid genotype classes (F_2_, B_1_, and B_2_), but with backcrosses to parental *T. m. triunguis* at the greatest frequency. Alabama and Florida GUxTT were only represented by *T. m. triunguis* backcrosses. Finally, *T. o. ornata* and *T. c. carolina* in Illinois displayed relatively few hybrid genotypes (5%) but all were F_1_, in contrast to the southeastern analyses (Fig. 2D).

### 3.3. Selective signatures at transcriptomic loci

Using Introgress, only SNPs with a high allelic frequency differential (*δ*>0.8) were retained (Andrés *et al.* 2013). One exception was the EAxGU analysis where *δ*>0.7 was applied because no loci at the higher threshold were recovered.

The Introgress genomic cline analysis recovered three outlier mRNA loci for EAxGU, and five each for EAxTT and GUxTT, with thirteen total and nine unique outliers among the three pairwise comparisons (Table 1; Fig. 3). Clines were inconsistent in that some pairwise taxon comparisons displayed rapid transitions from P_1_ to P_2_, whereas others demonstrated patterns indicative of directional introgression.

**Table 1:** Annotation information for overlapping Introgress and BGC (Bayesian Genomic Cline) outliers (Figs. 3, 4). Each outlier was derived from an alignment mapped to the *Terrapene* transcriptome, with three pairwise taxa combinations performed. The significance threshold (α_B_) was determined using a Bonferroni correction for multiple tests. Bold Gene abbreviations with an asterisk (*) differ significantly from neutral expectations. EA=Woodland (*T. c. carolina*), GU=Gulf Coast (*T. c. major*), TT=Three-toed (*T. m. triunguis*).

| Gene Abbr.<br>(P-value) | BGC ($\alpha/\beta$) | Outlier<br>($\alpha/\beta$)<sup>‡</sup> | Full Gene Name | Possible Function(s) | Source(s) |
| --- | --- | --- | --- | --- | --- |
| EAxGU ($\alpha_B=0.01$) | | | | | |
| **SULT (P=0)\*** | -1.59/-0.22 | -/N | Amine Sulfotransferase-like | Regulates yolk steroids during TSD<sup>†</sup> | (Paitz & Bowden 2008) |
| **TLR9 (P=0)\*** | -0.74/1.14 | -/N | Toll-like Receptor 9 | Immune Response to Pathogens | (Babik *et al.* 2015) |
| **ZNF236 (P=0)\*** | -0.94/0.14 | -/N | Zinc Finger Protein 236 | Unknown |  |
| EAxTT ($\alpha_B=0.007$) | | | | | |
| **SASH3 (P=0)\*** | -0.49/1.35 | N/+ | SAM and SH3 Domain Containing 3 | Immune response | (Kumaresan *et al.* 2018) |
| **SYPL2 (P=0)\*** | -0.68/1.65 | -/+ | Synaptophysin-like Protein 2 (AKA Mitsugumin 29) | Maintenance of [Ca<sup>2+</sup>] during anoxia | (Takeshima *et al.* 1998; Pamenter *et al.* 2016) |
| **FAM89B (P=0.036)** | -0.59/0.23 | -/N | Family with Sequence Similarity 89, member B | Upregulated in hypoxic conditions | (Goyal & Longo 2014) |
| **CITED4 (P=0)\*** | 0.26/0.97 | N/+ | Cbp/p300 Interacting Transactivator, Domain 4 | Inhibits hypoxia-activated transcription | (Fox *et al.* 2004) |
| **TLR9 (P=0)\*** | -0.38/1.63 | N/+ | Toll-like Receptor 9 | Immune Response to Pathogens | (Babik *et al.* 2015) |
| GUxTT ($\alpha_B=0.007$) | | | | | |
| **SASH3 (P=0)\*** | -0.22/1.86 | N/+ | SAM and SH3 Domain Containing 3 | Immune response | (Kumaresan *et al.* 2018) |
| **SYPL2 (P=0)\*** | -0.48/2.23 | -/+ | Synaptophysin-like Protein 2 | Maintenance of [Ca<sup>2+</sup>] during anoxia | (Takeshima *et al.* 1998; Pamenter *et al.* 2016) |
| **FAM89B (P=0)\*** | -0.46/0.66 | -/+ | Family with Sequence Similarity 89, member B | Upregulated in hypoxic conditions | (Goyal & Longo 2014) |
| **ACAD11 (P=0)\*** | -0.24/1.16 | N/+ | Acyl-CoA Dehydrogenase, family member 11-like | Lipid metabolism | (He *et al.* 2011; Gomez & Richards 2018) |
| **TMEM214 (P=0)\*** | 0.77/0.11 | +/N | Transmembrane Protein 214 | Stress-induced apoptosis during anoxia | (Kesaraju *et al.* 2009; Li *et al.* 2013) |

<sup>†</sup>TSD=Temperature-dependent sex determination
<sup>‡</sup> $\alpha$ outliers=excess P<sub>1</sub> (-) or P<sub>2</sub> (+) ancestry; $\beta$ outliers=slow (-) or rapid (+) genomic cline rates; N=neutral

**Figure 3:**
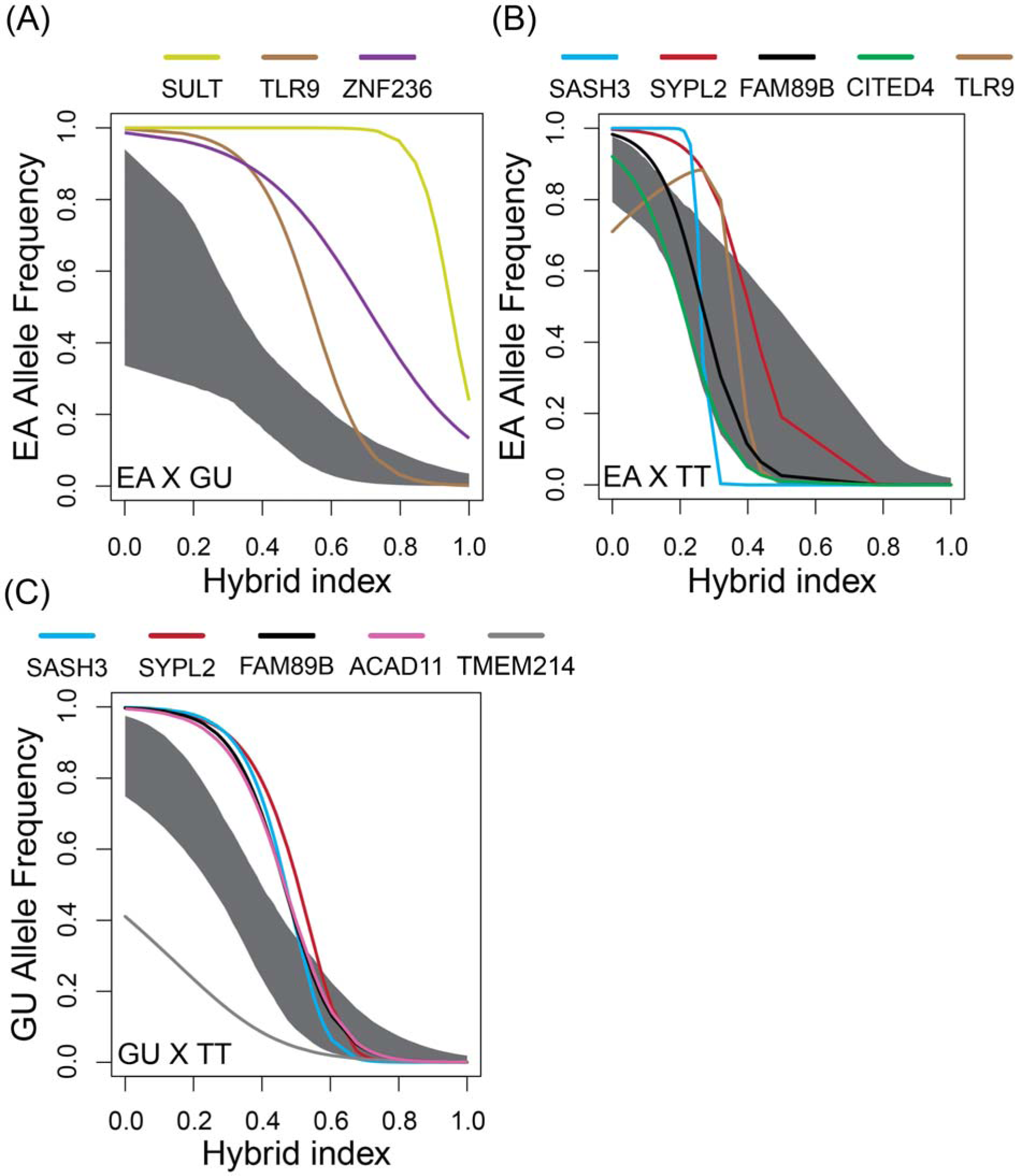
*Terrapene* Genomic clines depicting outlier SNPs found in transcriptome-aligned ddRAD loci. Pairwise comparisons are between EA=Woodland (*T. c. carolina*), GU=Gulf Coast (*T. c. major*), and TT=Three-toed (*T. mexicana triunguis*) box turtles. The gray area represents neutral expectations based on 2,660 (EAxGU), 2,623 (EAxTT), and 2,622 (GUxTT) transcriptome-aligned SNPs, and each line is a genomic cline for one outlier locus (abbreviations defined in Table 1).

For example, all three EAxGU outlier loci displayed an over-representation of EA alleles in the hybrid zone, concomitant with an under-representation of heterozygotes and GU alleles (Fig. 3A). The *SULT* locus was an extreme example, with excess EA alleles below a hybrid index of ~0.8 (=80% assignment to GU at diagnostic loci). This pattern was replicated to a lesser degree in *ZNF_2_36*, whereas *TLR9* was more sigmoidal. These genotypic proportions, coupled with the non-sigmoidal cline shape in *SULT* and *ZNF_2_36*, suggest that introgression may be driven by a directional shift towards homozygous P_1_ genotypes. In contrast, a steep, sigmoidal cline represented the genomic trend among scaffold assembly (and putatively non-functional) loci (Fig. 18). Taken together, these results suggest underlying directional introgression facilitating exchange of EA alleles despite divergence being maintained at most loci.

Cline shape was inconsistent within the EAxTT hybrid zone (Fig. 3B). Three (of five) outlier loci (*SASH3*, *SYPL2*, and *TLR9*) were significantly under-represented with regards to heterozygotes. Their clines displayed steep slopes, suggesting rapid transition among parental genotypes. An additional locus (*CITED4*) reflected an overrepresentation of P_2_ (TT), and a fifth (*FAM89B*) displayed three equally-represented genotypes. Of note, *FAM89B* did not differ significantly from neutral expectations following Bonferroni correction (*P*=0.036).

By contrast, neutral expectations were rejected in all five GUxTT clines (*P*=0; α=0.007), with four (i.e., *SASH3*, *SYPL2*, *ACAD11, FAM89B*) suggesting a pattern of restricted introgression (Fig. 3C). A fifth (i.e., *TMEM214*) displayed directional introgression, with the homozygous P_2_ (TT) genotype being overrepresented. Both the GUxTT and EAxTT analyses showed a ubiquitous signal of steep clines in non-transcriptomic loci (Fig. S18).

A greater number of SNP outliers (N=81) were identified with BGC rather than INTROGRESS, likely reflecting the larger number of loci included in the former. All INTROGRESS outliers were also corroborated by BGC (Fig. 4; Table 1). For EAxGU (Fig. 4A), all three exhibited excess EA ancestry (negative α=excess P_1_; positive α=excess P_2_), but there were no cline rate (ß) outliers in either direction. In contrast, four out of five EAxTT loci (*SYPL2*, *SASH3*, *TLR9*, and *CITED4*) were positive ß outliers, indicating steep clines and thus restricted introgression, with *SYPL2* also being an α outlier with excess EA (P_1_) ancestry (Fig. 4B). The fifth locus (*FAM89B*) was an α (not ß) outlier that favored EA alleles. Finally, GUxTT (Fig. 4C) included two loci that were both α and ß outliers with steep clines and excess GU ancestry (*SYPL2* and *FAM89B*). Two others were ß-only outliers with steep clines (*SASH3* and *ACAD11*), and one (*TMEM214*) an α-only outlier with excess TT ancestry.

**Figure 4:**
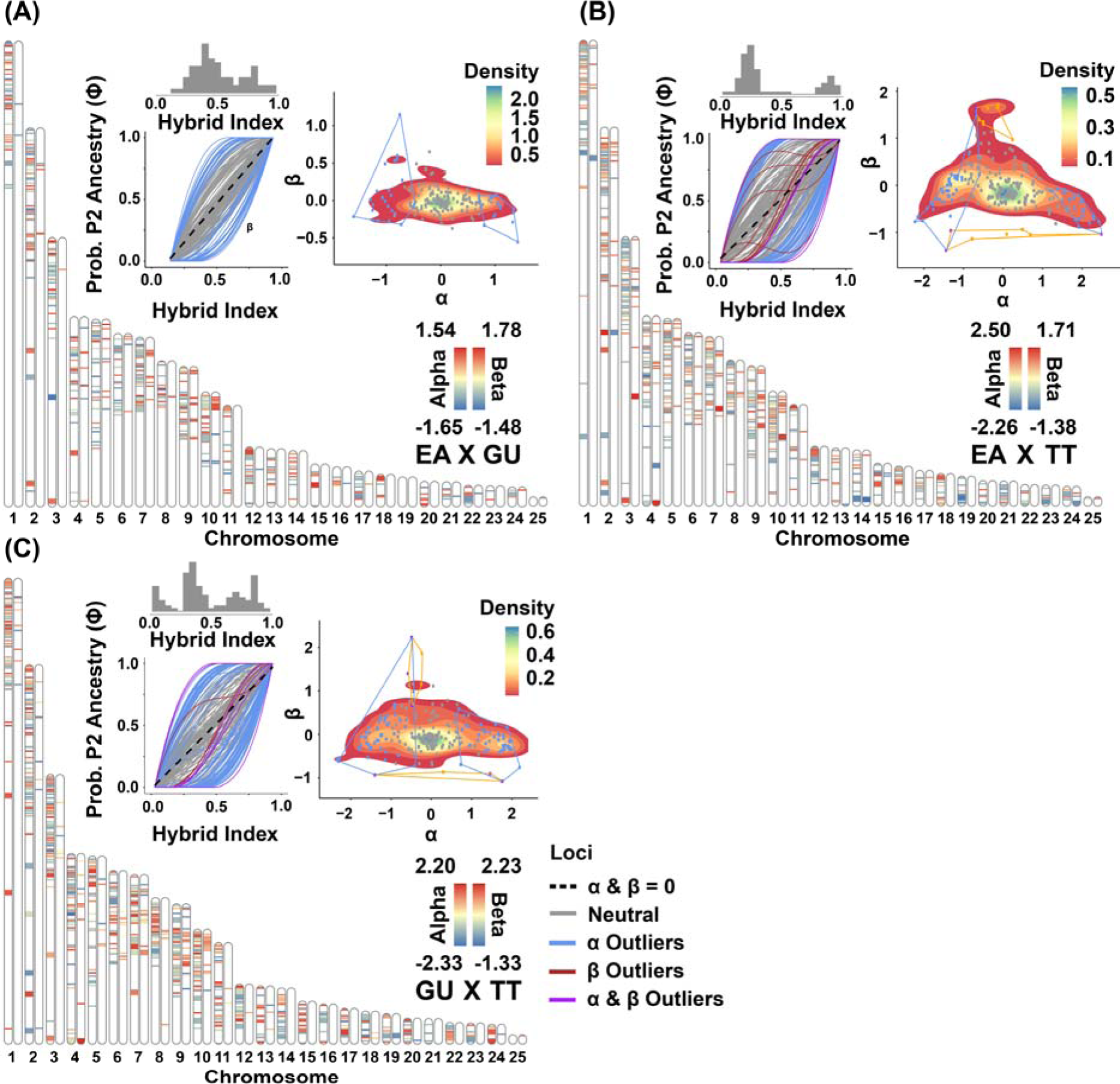
Bayesian genomic cline (BGC) outliers for *Terrapene* ddRADseq SNPs, plotted as heatmaps mapped to *Trachemys scripta* chromosomes. Each chromosome repeats to display significant outliers from the genomic cline center (α; left) and rate (ß; right) BGC parameters. Thinner and thicker bands represent SNPs from unknown scaffolds and annotated genes. BGC was run pairwise for three taxa: EA=*T. carolina carolina*, GU=*T. c. major*, and TT=*T. mexicana triunguis.* Outliers were significant if they had a 95% CI excluding zero or exceeding the probability distribution’s quantile interval 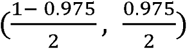. The plots depict transcriptome-aligned BGC genomic clines with hybrid indexs histogrsams above, and each line represents a genomic cline for one locus. The αXß plots illustrate the BGC parameters as a function of density. Polygons define density space for significant α(blue), ß (orange), and both (purple) outliers.

### 3.4 Environmental correlations with outliers

Shapiro-Wilks tests confirmed normality for all layers, following *OrderNorm* transformation. Forward selection retained ten uncorrelated and predictive layers (Table 2), and pRDAs revealed 49.2% environmental and 40.9% spatial contributions to model inertia. After controlling for spatial autocorrelation, the pRDA ANOVA identified four significant axes (*P*<0.05) explaining 21.5%, 10.6%, 10.3%, and 9.6% of the variance (cumulative 52.1%). The pRDA also identified 56 annotated outlier SNPs correlated with environmental predictors (Fig. 5). Twenty-eight pRDA outliers overlapped with Introgress and/or BGC analyses (Fig. 6). Many of the overlapping SNPs, including all nine Introgress loci that remained as outliers across all three analyses, were most strongly correlated with temperature variables rather than precipitation or wind speed (Table 3).

**Table 2:** WorldClim environmental predictor abbreviations for the redundancy analysis (RDA). The raster layers were obtained from https://worldclim.org. Additional variables were excluded via forward selection due to low predictive capacity or correlation with the remaining layers.

| Abbr. | Full Name | BioClim No. |
| --- | --- | --- |
| tmDR2 | Mean diurnal range [Monthly (Max temp - Min temp)] | 2 |
| tS4 | Temperature seasonality (standard deviation X 100) | 4 |
| mxtWM5 | Max temperature of warmest month | 5 |
| mintCM6 | Min temperature coldest month | 6 |
| mtWQ8 | Mean temperature of wettest quarter | 8 |
| pAM12 | Annual precipitation | 12 |
| pDM14 | Precipitation of driest month | 14 |
| pWQ16 | Precipitation of wettest quarter | 16 |
| pCQ19 | Precipitation of coldest quarter | 19 |
| Wind | Mean annual wind speed | N/A |

**Table 3:** *Terrapene* ddRAD SNP outliers (N=118) most strongly associated with ten predictive and uncorrelated WorldClim variables (Table 2), collapsed into temperature, precipitation, or wind speed categories. Percentages include the total for only redundancy analysis (% RDA) and for all three outlier methods (% All): Introgress, Bayesian genomic cline (BGC), and RDA.

| **Environment**<br>**Type** | **No.**<br>**Outliers** | **%**<br>**(RDA)** | **%**<br>**(All)** |
| --- | --- | --- | --- |
| Temperature | 44 | 78.6 | 34.7 |
| Precipitation | 10 | 17.9 | 8.5 |
| Wind | 2 | 3.6 | 1.7 |
| Total | 56 | 100.0 | 44.9 |

**Figure 5:**
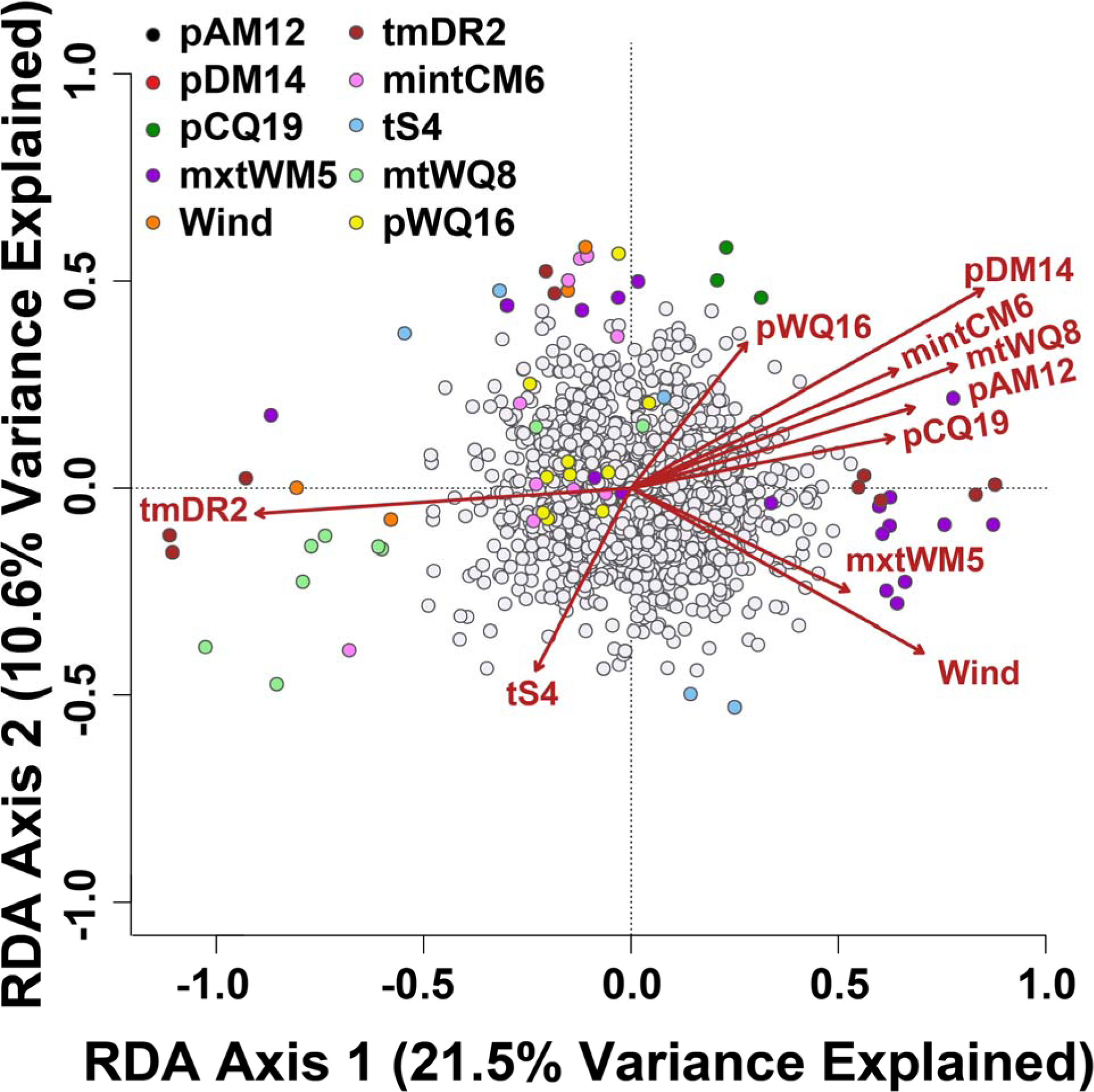
Redundancy analysis (RDA) representing outlier *Terrapene* SNPs correlated with ten predictive, non-redundant BioClim environmental variables (see Table 2 for predictor abbreviations). Significant outliers were designated as being +/− 3 standard deviations from the RDA axis loading means. Pairwise Pearson’s correlations between each outlier and environmental variable were performed, and the strongest correlation coefficient (r) determined the best-supported predictor.

**Figure 6:**
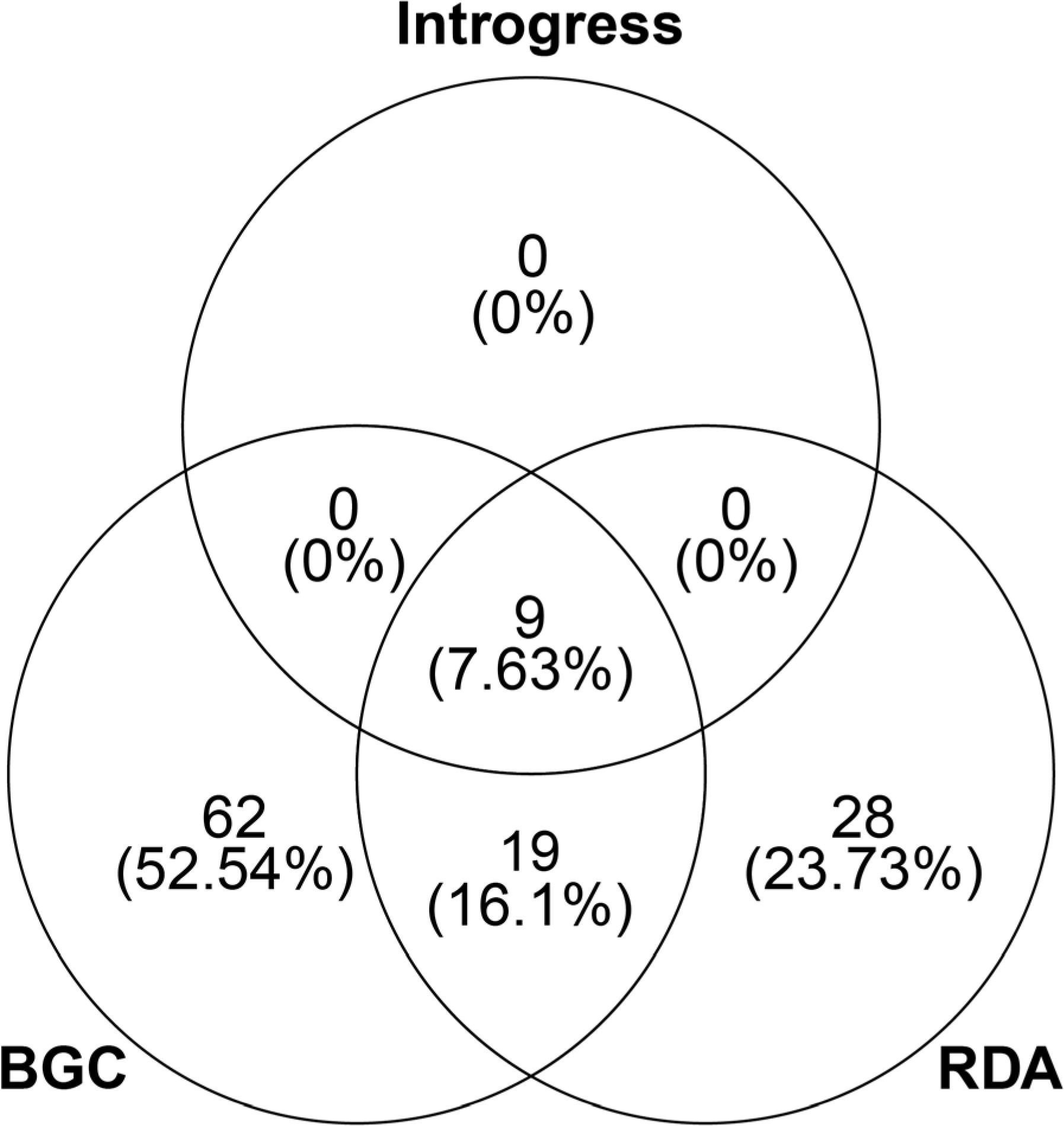
Venn Diagram depicting overlap between *Terrapene* transcriptomic outliers identified in INTROGRESS, Bayesian genomic cline (BGC), and redundancy analysis (RDA). Each value includes raw counts (top) and percentages (bottom).

## 4. DISCUSSION

Our analyses characterized introgression in two North American box turtle hybrid zones (i.e., midwestern and southeastern North America). The midwestern hybrid zone showed no evidence of introgression, with hybrids restricted at low frequency to F_1_, whereas southeastern hybridization was introgressive in nature, as evidenced by numerous backcrosses and F_2_ individuals and a conspicuous lack of F_1_ hybrids. Furthermore, contrasting intra- and inter-specific southeastern hybrid zones revealed they not only varied in the genealogical composition of their hybrids but also in the shapes and widths of locus-specific clines. We propose such a contrast provides insight into the evolutionary histories of the taxa involved and serves to delineate their appropriate taxonomic designations. Specifically, recent phylogenetic research indicates that *T. m. triunguis* is a separate species from *T. c. carolina* and *T. c. major* (Martin *et al.* 2013, 2014, 2020). The genomic clines herein are consistent with those phylogenetic results in that transcriptomic loci with steep clines are only found in inter-specific comparisons (i.e., *T. mexicana versus T. carolina*). The candidate genes at these transcriptomic loci may be targets of selection and directional introgression, as they deviated from neutral expectations not only with respect to genome-wide ancestry (i.e., genomic clines) but also multivariate environmental associations. Below we consider the impact of these results on the evolutionary history, genomic architecture, and species boundaries of *Terrapene*.

### 4.1. Regional and Taxon-specific Perspectives

Several of our conclusions have implications for *Terrapene* systematics. First, our Admixture analyses substantiate the presence of discrete *T. c. major* populations in Florida and Mississippi, with the Alabama and Apalachicola river drainages potentially serving as biogeographic barriers (Fig. S8). This consideration was markedly absent in previous morphological analyses (Butler *et al.* 2011) that concluded *T. c. major* merely represented an area of Admixture between other *Terrapene* in the region. We did indeed detect considerable Admixture between *T. c. major*, *T. c. carolina*, and *T. m. triunguis*, but the presence of two apparently non-admixed *T. c. major* populations demonstrated the existence of cryptic genetic variation (Douglas *et al.* 2009). We interpret these populations as representing distinct evolutionary significant units (ESUs) or (at worst) management units (MUs).

Second, Admixture is apparent among all southeastern *Terrapene* except *T. bauri*, which absorbed the greatest amount of DAPC variation. This is likely attributed to Pliocene vicariance in the Florida Peninsula, where the Okefenokee Trough divided northern from southern Florida (Bert 1986; Douglas *et al.* 2009). Each of these aspects will require careful consideration when conservation efforts are planned or implemented, particularly given that *Terrapene* are in decline throughout their range (Dodd 2001).

On the other hand, *T. ornata* and *T. carolina* are separated by greater genetic distances than are the southeastern taxa (Martin *et al.* 2013), which may suggest the presence of intrinsic genetic incompatibilities (Barton 2001; Abbott *et al.* 2013) and is consistent with the lack of hybrids beyond the F_1_ generation. Furthermore, the low frequency of F_1_ hybrids observed in the Illinois ONxEA population may have resulted from recent degradation of the prairie grassland habitat (Manning 2001; Mussmann *et al.* 2017), which subsequently initiated increased heterospecific contact or otherwise disturbed reproductive boundaries by altering the fitness consequences of hybridization (Chafin et al. 2019; Grabenstein and Taylor 2018). However, this hypothesis cannot be explicitly tested herein. Although we did not identify contemporary back-crossed individuals, Cureton *et al.* (2011) did depict potential introgressive hybridization between ONxEA based on mitochondrial DNA and multiple microsatellite markers. These discrepancies may reflect either historical Admixture as a source of introgression, which we did not explicitly test for, or back-crossed hybrids as a rarity not encountered in our sampling.

### 4.2. Biogeography and hybrid zone formation in Terrapene

The disparity in early and late-generation hybrids between the midwestern (ONxEA) and southeastern (EAxGU, GUxTT, and EAxTT) hybrid zones suggests differences in the underlying evolutionary processes. Such differences could involve regional variability in the extent or nature of reproductive isolation, or simply their respective biogeographic histories. Pleistocene glaciation precipitated numerous widespread distributional shifts across many taxa in the Midwest (King 1981; Webb 1981). Subsequent postglacial “shuffling” has also been implicated in as an historical driver of introgressive hybridization in populations of *Sistrurus* rattlesnakes now allopatric in the same region (Sovic *et al.* 2016). The same process could explain evidence for past introgression in the ONxEA hybrid zone (Cureton *et al* 2011b), despite a lack of contemporary hybrids. Here, a rapid re-colonization [especially from northern refugia such as the “Driftless Area” of Wisconsin, Iowa, and Illinois during the last glacial maximum (Holliday *et al.* 2002)] resulted in north-south contact zones near the glacial maximum (i.e., “leading-edge” hypothesis; Hewitt 1996, 2000). However, our purported range overlap also represents a broad interdigitation of “prairie” and “interior highland” habitats (Ennen *et al.* 2017), indicating that later-generation hybrids simply requires a finer-scale sampling than ours to be detected.

The southeastern United States has long been known for the of co-occurrence of contact zones (Remington 1968; Avise 2000), with migration from refugia in southern Florida and east Texas/ west Louisiana as an hypothesized mechanism (Swenson & Howard 2005). The divergence of these lineages prior to Pleistocene glaciation (Martin *et al.* 2013) seemingly indicates postglacial expansion as a potential mechanism underlying southeastern hybrid zone formation. Likewise, finer-scale phylogeographic breaks that corroborate those in *Terrapene* have also been detected in numerous other turtles (Walker & Avise 1998). For example, *Sternotherus minor* and *S. odoratus* show deep east-west phylogeographic breaks approximately centered on Alabama, with unique lineages in peninsular Florida (Iverson 1977; Walker & Avise 1998). *Kinosternon subrubrum* mirrors this pattern, but with an additional unique lineage in the panhandle region (Walker *et al.* 1998). Similar breaks again appear in *Trachemys scripta, Macroclemys temminckii* (Walker & Avise 1998; Roman *et al.* 1999), and *Gopherus polyphemus* (Lamb *et al.* 1989). It is thus no surprise for the region to be identified as a hotspot for inter-specific contact and phylogeographic concordance (Soltis *et al.* 2006; Rissler & Smith 2010), also reflected in *Terrapene*. As always, it becomes inherently difficult to separate the relative importance of historical processes from contemporary physiographic or ecological features.

### 4.3. Functional Genomic Architecture in the Southeastern Hybrid Zone

Regardless of the biogeographic scenarios invoked, a clear hotspot for hybridization exists in the southeastern United States. The variance among locus-specific patterns of differentiation or exchange allows for the potential interpretation of adaptive processes, at least in the context of neutral expectations. SNPs located in several mRNA loci are implicated as potentially contributing to selection in *Terrapene* from three southeastern hybrid zones. Among inter-specific comparisons (Figs. 3B, 3C, 4B, 4C), the dominant pattern was a steep, sigmoidal cline (GUxTT and EAxTT, but most clearly apparent in the latter). This accordingly points to selection against interspecific heterozygotes (Fitzpatrick 2013). In contrast, the selective advantage of EA alleles in the EAxGU hybrid zone (Figs. 3A, 4A) fails to agree with the general genome-wide pattern of underdominance (Fig. S18), which suggests directional introgression within which EA alleles are favored in hybrids under contemporary conditions. Mapping BGC outliers to *Trachemys* chromosomes (Fig. 4) also revealed their ubiquitous rather than finely concentrated distribution across the genome, and this in turn highlighted the differential nature of the observed interspecific introgression. Inaccuracies regarding mapping a *Terrapene* assembly against a *Trachemys* reference genome is also a possibility, although we attempted to minimize this by resolving conflicts via PAFScaff.

Loci with steep genomic clines were most strongly correlated with temperature predictors, suggesting the importance of thermal adaptations in maintaining species boundaries between southeastern *Terrapene*. In contrast, neither precipitation nor wind-associated outliers followed this pattern. Given the positive relationship between outlier genes and thermal predictors, a natural extrapolation would be that a thermal gradient drives differential introgression. This has multiple implications regarding the integrity of species boundaries if *Terrapene* are subjected to future environmental changes. In one scenario, a shifting adaptive landscape may promote hybridization by contravening long-term reproductive isolation (EAxGU), with subsequent introgression at specific loci (as herein). Alternatively, rapid environmental change could simply outpace the selective filtering of maladaptive variants, with a subsequent decrease in fitness (Kokko *et al.* 2017). This would be particularly evident when effective population sizes are already depressed following a population bottleneck (Chafin *et al.* 2019). Here, extreme rates of change may also link with a genetic swamping effect (i.e., replacement of local genotypes by hybrids; Todesco *et al.* 2016). Both scenarios implicate anthropogenic pressures as governing the fates of diverse taxa across hybrid zones (Taylor *et al.* 2015).

The putative functions of the nine outlier loci supported by genomic clines and RDA provide additional support for the strong impact of thermal selection. Two loci potentially relate to TSD during embryonic development, while others seemingly associate with molecular pathways in skeletal muscle and nervous tissues that involve tolerance to anoxia and hypoxia (N=6), and immune response to pathogens [(N=2); see Table 1 for sources]. Anoxia/ hypoxia-related genes have been associated with freeze tolerance in hibernating turtles (Storey 2006), thus supporting an obvious association with thermal gradients. Here, three loci (*SYPL2, ACAD11*, and *TMEM214*) may regulate brain function and metabolism by up-regulating Ca^2+^ concentrations (Takeshima *et al.* 1998; Pamenter *et al.* 2016), inducing lipid metabolism (He *et al.* 2011; Gomez & Richards 2018), and initiating stress-induced apoptosis (Kesaraju *et al.* 2009; Li *et al.* 2013). Similarly, the *CITED4* gene (EAxTT) potentially inhibits hypoxia-related transcription factors (Fox *et al.* 2004), whereas *FAM89B* (GUxTT) may become up-regulated when physiological conditions turn hypoxic (Goyal & Longo 2014). The regulation of immune function is less clearly associated with underlying thermal gradients, but may associate instead with behavioral thermoregulation during infection, given that infection resistance increases at warmer temperatures (Dodd 2001; Agha *et al.* 2017). However, we remain cautious in that these genes have not been associated with specific functions in *Terrapene*, and to do so remains speculative. Nevertheless, their potential connection with thermal adaptations is consistent with the RDA.

We emphasize that ectothermic vertebrates are exceptionally vulnerable to contemporary pressures, and reflect an elevated extinction risk due to a strong reliance on environmental thermoregulation and a dependence on suitable habitat (Gibbons *et al.* 2000; Sinervo *et al.* 2010; Winter *et al.* 2016). Indeed, ectotherms can exhibit reduced fitness and growth-rates when environmental conditions exceed their thermal optima (Deutsch *et al.* 2008; Martin & Huey 2008; McCallum *et al.* 2009; Huey *et al.* 2012; Huey & Kingsolver 2019). Increased temperatures also impact physiological pathways, and these are putatively comparable with the genes described herein, such as increasing metabolic rates (Dillon *et al.* 2010), intensifying hypoxic stress despite higher temperature-driven O_2_ demands (Huey & Ward 2005), heightening disease transmission (Pounds *et al.* 2006), or even the over-extension of thermal tolerances (Sinervo *et al.* 2010; Ceia-Hasse *et al.* 2014). Going forward, climate change may facilitate evolutionary responses to changing thermal conditions, potentially including local adaptation (Holt 1990; Norberg *et al.* 2012; Bush *et al.* 2016), physiological and behavioral mechanisms (e.g., thermoregulation, phenology), plasticity (Urban *et al.* 2014; Sgrò *et al.* 2016), of shifts in species distributions (Parmesan *et al.* 1999; Parmesan & Yohe 2003; Moreno-Rueda *et al.* 2012). Accordingly, ectothermic species boundaries may be particularly susceptible if indeed governed by thermal conditions, as seemingly exemplified herein with *Terrapene*.

### 4.3 Conclusions

Our study suggests that reproductive isolation in turtles involves numerous mechanisms regulated by thermally induced selective pressures. Clearly, such selective pressures play a prominent role in chelonian ecology. Many turtle species are at elevated risks from climate change due to the imposition of TSD during embryonic development, as well as prolonged generation times. Similarly, long generation times can also restrict the adaptive capacity of turtles in a rapidly changing climate (Hoffmann *et al.* 2017). Here, climate change can shift sex-ratios and promote demographic collapse, with warmer temperatures initiating male bias in biological sex, and vice versa (Janzen 1994).

Two important evolutionary implications are evident in our data. First, we demonstrated differential introgression along an ecological gradient in three taxa inhabiting a North American hybrid zone. We then assessed scaffold-aligned and transcriptomic SNPs to identify several genes whose functions are consistent with physiological processes related to thermal ecology, and as such are capable of promoting adaptive divergence. In this sense, they potentially describe ecological gradients related to TSD, anoxia/ hypoxia tolerance, and immune response in a hybrid zone encompassing three taxa. While we acknowledge that the observed clinal patterns could represent “molecular spandrels” reflecting an underlying neutral process, such as isolation-by-distance, we sought environmental associations by actively controlling for spatial autocorrelation as a corroboration of clinal loci with a putative adaptive role (Vasemägi 2006; Barrett & Hoekstra 2011). We also underscore specific loci displaying genomic cline patterns consistent with directional introgression and selection, and these loci are potentially sustaining divergence across species.

Second, we characterized a southeastern North American hybrid zone representing a variety of biodiversity elements as being susceptible to anthropogenic and environmental changes (Remington 1968; Swenson & Howard 2005; Rissler & Smith 2010). Our results demonstrate that NGS population genomic methods can clearly identify population structure and detect introgression, whereas traditional Sanger sequencing methods are inadequate to do so (Butler *et al.* 2011; Martin *et al.* 2013). We also underscore specific loci prone to directional introgression and selection that may potentially sustain divergence across species.

## ACKNOWLEDGEMENTS

The research herein was conducted in partial fulfillment of the Ph.D. degree in Biological Sciences at the University of Arkansas (BTM). We would like to extend our gratitude to the countless volunteers and organizations who collected and/or provided tissue samples (Table S1). We also thank the current and former members of the Douglas Lab and University of Arkansas faculty who provided advice and support, especially A. Alverson, W. Anthonysamy, M. Bangs, J. Koukl, S. Mussmann, J. Pummill, and Z. Zbinden. Finally, many thanks to the reviewers, whose comments led to a greatly improved manuscript. Sample collections were approved under the University of Texas-Tyler Animal Care and Use Committee (IACUC) permit #113 and University of Illinois IACUC protocols 16160 and 18000. Funding was provided by the Lucille F. Stickle Fund of the North American Box Turtle Committee, the American Turtle Observatory (ATO), and the following endowments: The Bruker Professorship in Life Sciences (MRD) and the Twenty-First Century Chair in Global Change Biology (MED). Analytical resources were provided by the Arkansas High Performance Computing Cluster (AHPCC) and an NSF-XSEDE Research Allocation (TG-BIO160065) that allowed access the Jetstream cloud service.

## DATA ACCESSIBILITY

The demultiplexed ddRADseq reads are deposited as FASTQ files to NCBI’s sequence read archive (https://www.ncbi.nlm.nih.gov/sra); Accessions: SAMN12668545-SAMN12668981 (BioProject ID: PRJNA563121). The R scripts, metadata, and input files for each analysis are available from a Dryad Digital Repository (https://doi.org/10.5061/dryad.brv15dv7k).

## AUTHOR CONTRIBUTIONS

BTM and TKC conceived the research, laboratory, and analytical tools, scripts, and approaches. BTM performed the lab work, analyzed the data, conducted bioinformatic analyses, and wrote the manuscript. MRD and MED were the study supervisors, guided the study design, and provided funding. JSP facilitated the collection of thousands of *Terrapene* tissues and provided expertise in methodological development. RDB collected hundreds of *Terrapene* tissues from southeastern North America and facilitated the collection of many additional individuals. CAP provided tissues from the midwestern hybrid zone as well as sample site expertise in Illinois. All authors contributed to editing and revising the manuscript

