## Supplementary Tables / Figures for "Contrasting signatures of introgression in North American box turtle (*Terrapene* spp.) contact zones"

**Table of Contents:**

| **Table S1** | Separate File |
| --- | --- |
| **Table S2** | Page 2 |
| **Table S3** | Page 3 |
| **Figure S1** | Page 4 |
| **Figure S2** | Page 5 |
| **Figure S3** | Page 6 |
| **Figure S4** | Page 7 |
| **Figure S5** | Page 8 |
| **Figure S6** | Page 9 |
| **Figure S7** | Page 10 |
| **Figure S8** | Page 11 |
| **Figure S9** | Page 12 |
| **Figure S10** | Page 13 |
| **Figure S11** | Page 14 |
| **Figure S12** | Page 15 |
| **Figure S13** | Page 16 |
| **Figure S14** | Page 17 |
| **Figure S15** | Page 18 |
| **Figure S16** | Page 19 |
| **Figure S17** | Page 20 |
| **Figure S18** | Page 21 |

**Table S2:** Number of sequenced individuals (N) per *Terrapene* taxon, as identified in the field. *T. carolina carolina*=Woodland, *T. c. major*=Gulf Coast, *T. c. bauri*=Florida, *T. carolina*=field identification limited to species-level, *T. m. triunguis*=Three-toed, and *T. o. ornata*=Ornate box turtles.

| **Taxonomic ID** | **N** |
| --- | --- |
| *T. carolina carolina* | 106 |
| *T. carolina major* | 88 |
| *T. carolina bauri* | 4 |
| *T. carolina* | 65 |
| *T. mexicana triunguis* | 47 |
| *T. ornata ornata* | 62 |
| **Total** | 368 |

**Table S3:** Genotype frequency proportions from four NewHybrids analyses involving the GU=Gulf Coast (*T. c.* *major*), EA=Woodland (*T. c.* *carolina*), TT=Three-toed (*T. m. triunguis*), and ON=Ornate (*T. o. ornata*) box turtles; TC=*T. carolina* (subspecies unidentified). The second two letters in the population ID correspond to U.S. state locality (AL=Alabama, FL=Florida, LA=Louisiana, SC=South Carolina, GA=Georgia, MS=Mississippi, IL=Illinois). Columns depict the proportion of assignment to parental (P_1_ and P_2_), first and second-generation hybrid (F_1_ and F_2_), backcross (B_1_ and B_2_), and unassigned (FN) genotype frequency classes.

| **Population** | | **P1** | **P2** | **F1** | **F2** | **B1** | **B2** | **FN** |
| --- | --- | --- | --- | --- | --- | --- | --- | --- |
| **GUxEA** | |  |  |  |  |  |  |  |
|  | PureGU | 1.00 | 0.00 | 0.00 | 0.00 | 0.00 | 0.00 | 0.00 |
|  | PureEA | 0.00 | 1.00 | 0.00 | 0.00 | 0.00 | 0.00 | 0.00 |
|  | EAAL | 0.00 | 1.00 | 0.00 | 0.00 | 0.00 | 0.00 | 0.00 |
|  | GUFL | 0.46 | 0.04 | 0.00 | 0.08 | 0.21 | 0.00 | 0.21 |
|  | TCAL | 0.02 | 0.86 | 0.00 | 0.02 | 0.02 | 0.00 | 0.08 |
|  | GUAL | 0.20 | 0.20 | 0.00 | 0.00 | 0.00 | 0.00 | 0.60 |
| **EAxTT** | |  |  |  |  |  |  |  |
|  | PureEA | 1.00 | 0.00 | 0.00 | 0.00 | 0.00 | 0.00 | 0.00 |
|  | PureTT | 0.00 | 1.00 | 0.00 | 0.00 | 0.00 | 0.00 | 0.00 |
|  | TTLA | 0.00 | 1.00 | 0.00 | 0.00 | 0.00 | 0.00 | 0.00 |
|  | EASC | 0.47 | 0.00 | 0.00 | 0.00 | 0.40 | 0.00 | 0.13 |
|  | TCGA | 0.80 | 0.00 | 0.00 | 0.00 | 0.00 | 0.10 | 0.10 |
|  | EAGA | 0.91 | 0.00 | 0.00 | 0.00 | 0.05 | 0.00 | 0.05 |
|  | TCAL | 0.96 | 0.00 | 0.00 | 0.00 | 0.04 | 0.00 | 0.00 |
| **TTxGU** | |  |  |  |  |  |  |  |
|  | PureTT | 0.00 | 1.00 | 0.00 | 0.00 | 0.00 | 0.00 | 0.00 |
|  | PureGU | 1.00 | 0.00 | 0.00 | 0.00 | 0.00 | 0.00 | 0.00 |
|  | GUAL | 0.60 | 0.00 | 0.00 | 0.00 | 0.20 | 0.00 | 0.20 |
|  | TTMS | 0.00 | 0.50 | 0.00 | 0.17 | 0.00 | 0.06 | 0.28 |
|  | TTLA | 0.00 | 1.00 | 0.00 | 0.00 | 0.00 | 0.00 | 0.00 |
|  | TCMS | 0.43 | 0.00 | 0.00 | 0.00 | 0.57 | 0.00 | 0.00 |
|  | GUMS | 0.52 | 0.02 | 0.00 | 0.00 | 0.13 | 0.04 | 0.28 |
|  | GUFL | 0.88 | 0.04 | 0.00 | 0.00 | 0.04 | 0.00 | 0.04 |
| **ONxEA** | |  |  |  |  |  |  |  |
|  | PureON | 1.00 | 0.00 | 0.00 | 0.00 | 0.00 | 0.00 | 0.00 |
|  | PureEA | 0.00 | 1.00 | 0.00 | 0.00 | 0.00 | 0.00 | 0.00 |
|  | ONIL | 0.74 | 0.21 | 0.05 | 0.00 | 0.00 | 0.00 | 0.00 |
|  | EAIL | 0.00 | 0.98 | 0.00 | 0.00 | 0.00 | 0.00 | 0.03 |

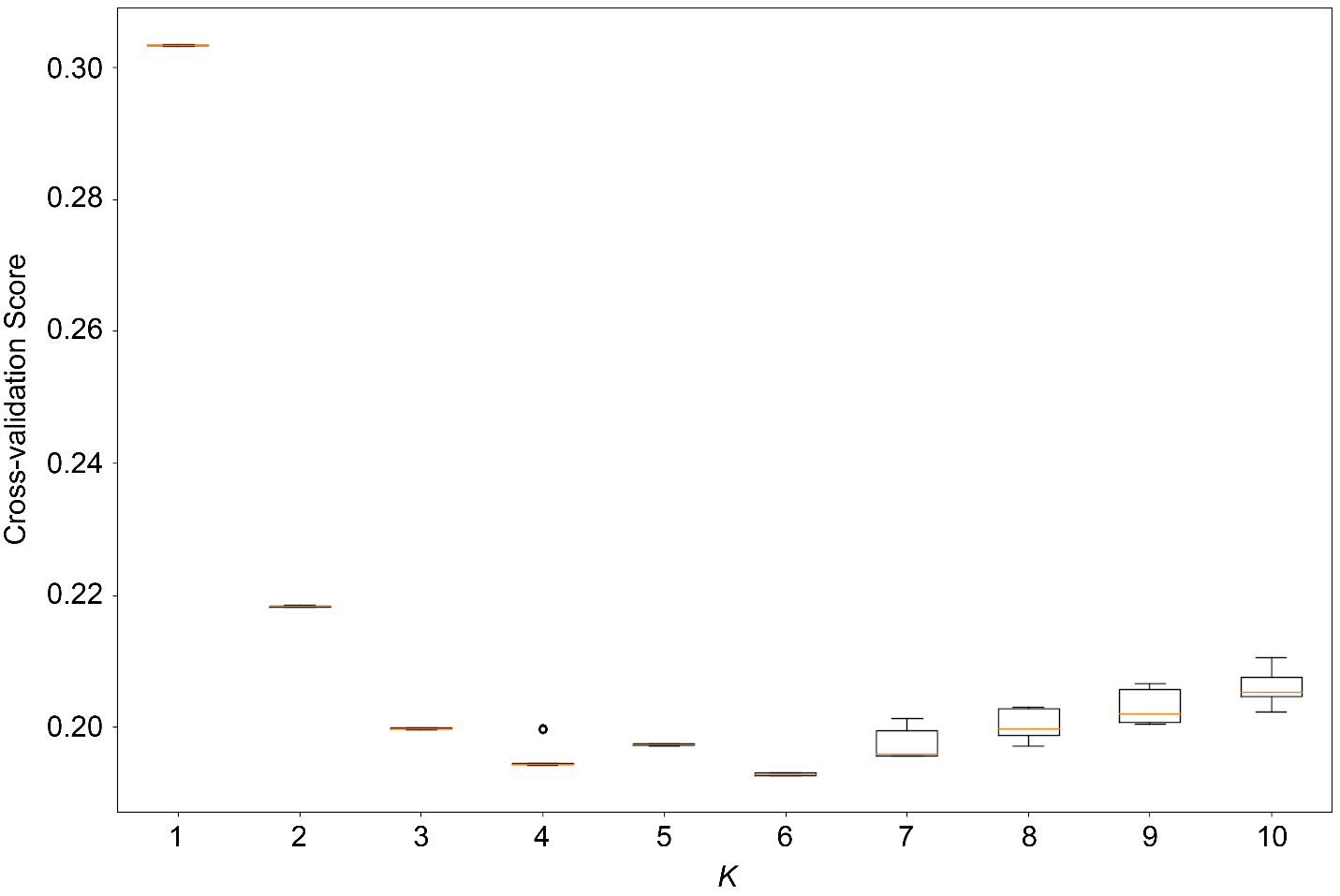

**Figure S1:** Cross-validation (CV) scores across *K-values* (*K*=1-10) for Admixture runs containing all sequenced samples (N=368). Lower CV values indicate less error and stronger support for the corresponding *K.*

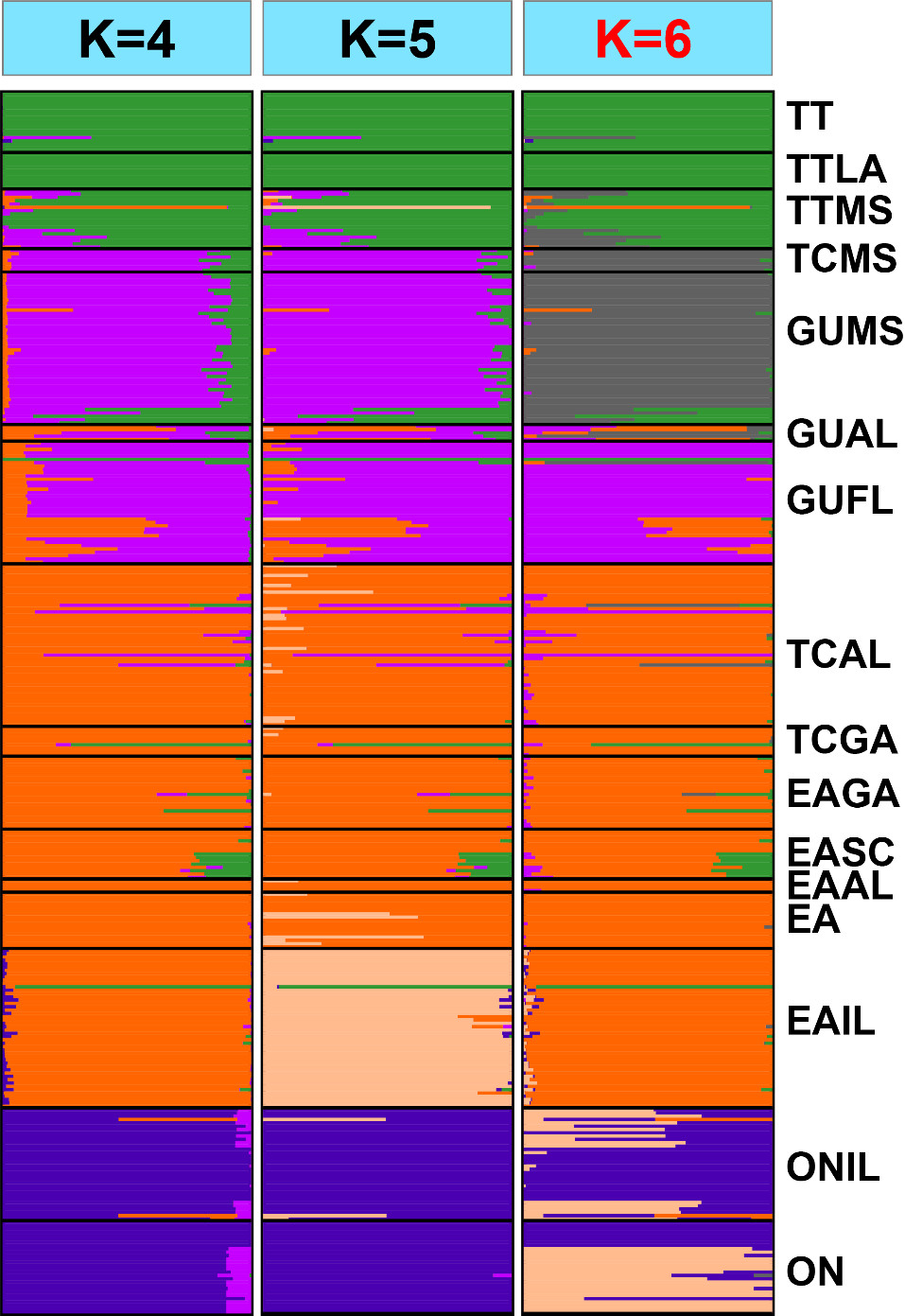

**Figure 1:** Top three *Terrapene* Admixture plots representing 12,052 unlinked ddRAD SNPs across all sampled populations. The lowest cross-validation score was for *K*=6 (depicted at right in red), followed by *K*=4 and then *K*=5. Each bar represents a unique individual, and bars with mixed colors represent admixed ancestry. The first two letters of the populations correspond to subspecific field identification (ON=Ornate, *T. ornata ornata*; EA=Woodland, *T. carolina carolina*; GU=Gulf Coast, *T. c. major*; TT=Three-toed, *T. mexicana triunguis*; TC=*Terrapene carolina*, with subspecies unidentified in the field). The second two letters (if present) represent locality codes for U.S. or Mexican state (IL=Illinois; AL=Alabama; GA=Georgia; SC=South Carolina; FL=Florida; MS=Mississippi; LA=Louisiana). Populations lacking a state locality code consisted of multiple localities sampled outside hybrid zones.

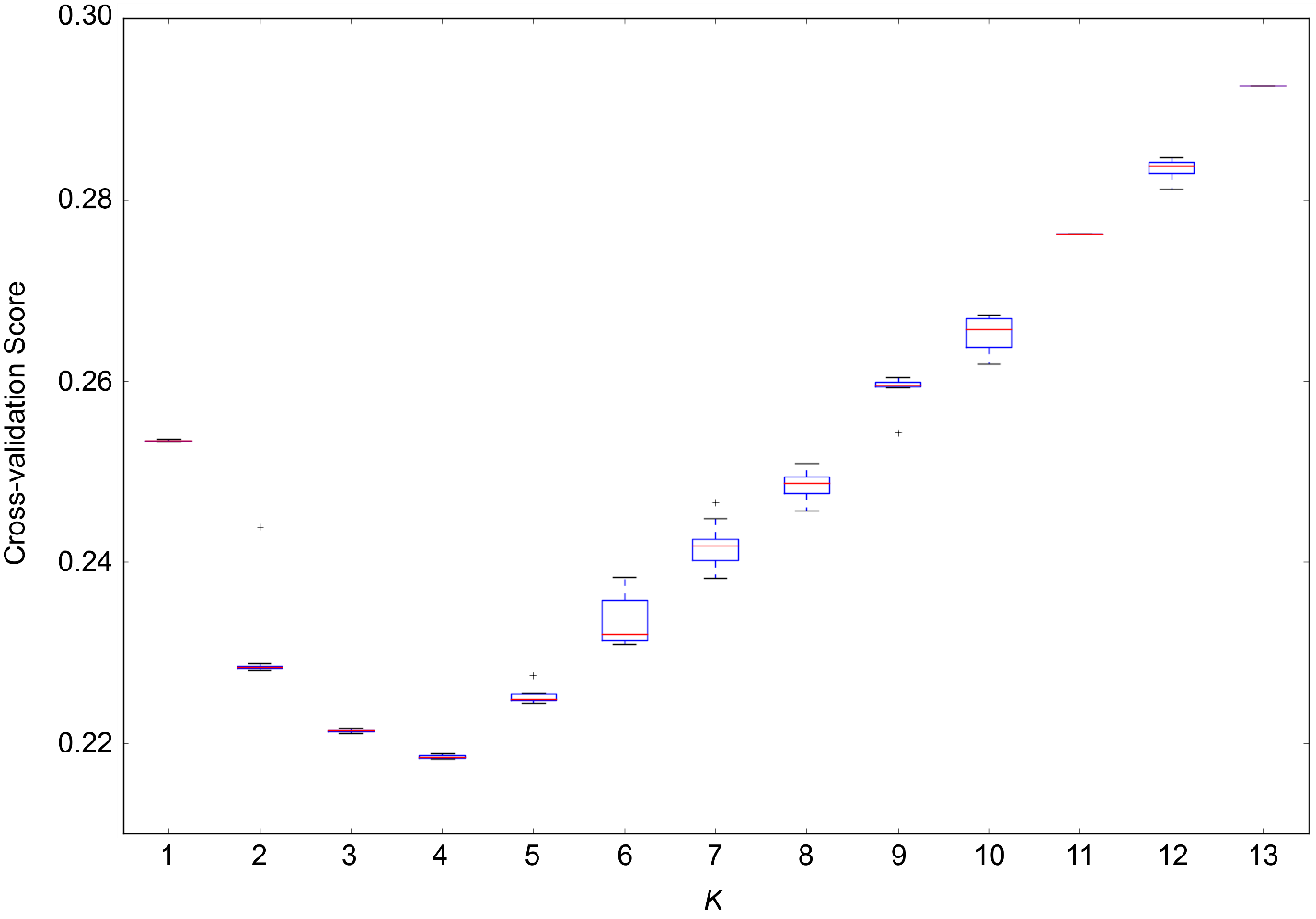

**Figure S3**: Cross-validation (CV) scores across *K-values* (*K*=1-13) for Admixture runs containing samples from southeastern North America (N=259). Lower CV values indicate less error and stronger support for the corresponding *K*.

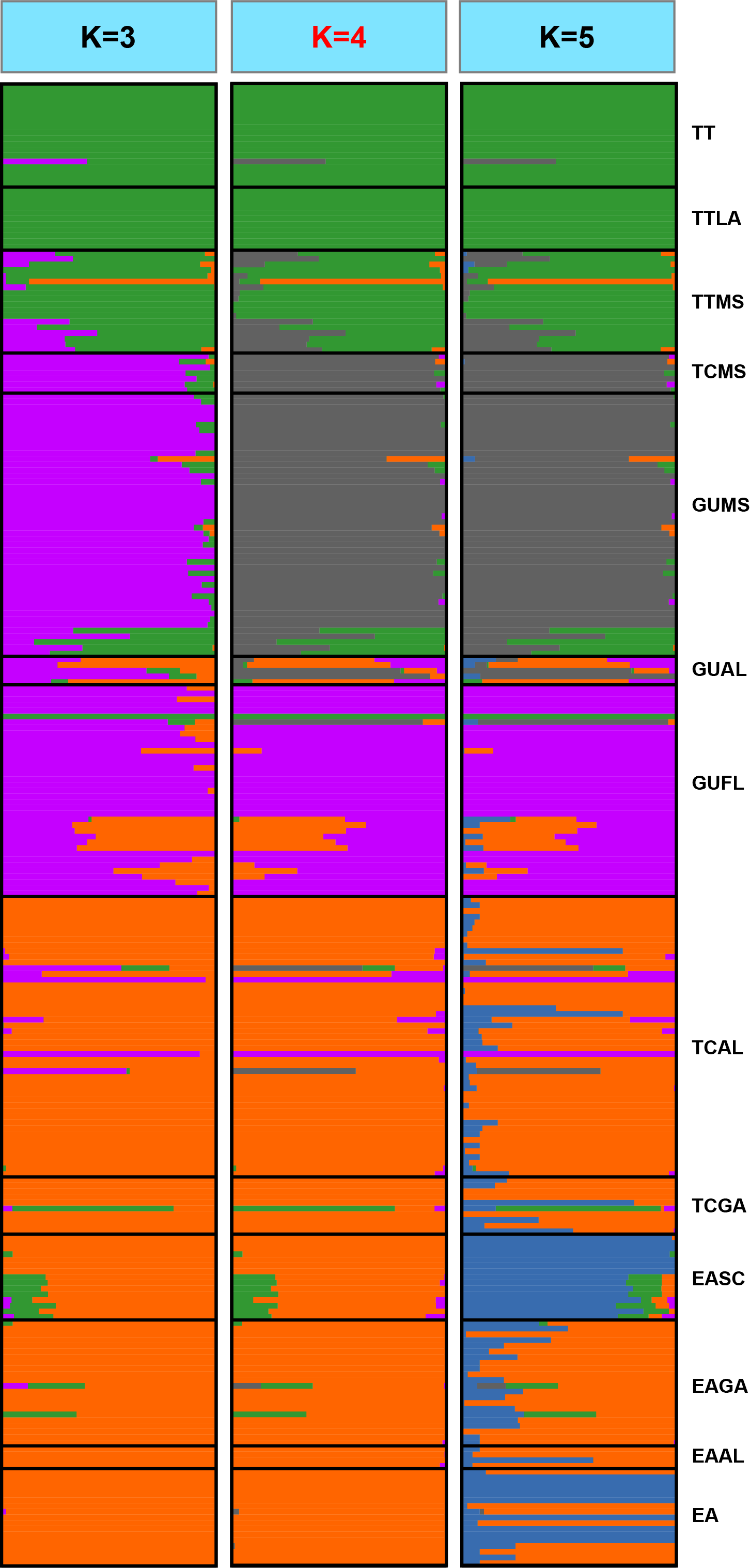

**Figure S4:** Top three southeastern *Terrapene* Admixture plots representing 11,308 unlinked ddRAD SNPs. The lowest cross-validation score was for *K*=4 (depicted in red), followed by *K*=3 then *K*=5. Each bar represents a unique individual, and bars with mixed colors depict admixed ancestry. The first two population code letters correspond to subspecific field identification (EA=Woodland, *T. c. carolina*; GU=Gulf Coast, *T. c. major*; TT=Three-toed, *T. m. triunguis*; TC=*Terrapene carolina*, with subspecies unidentified). The second two letters represent locality codes for U.S. states (AL=Alabama; GA=Georgia; SC=South Carolina;FL=Florida; MS=Mississippi; LA=Louisiana). Populations lacking a state code consisted of multiple localities sampled outside the hybrid zone.

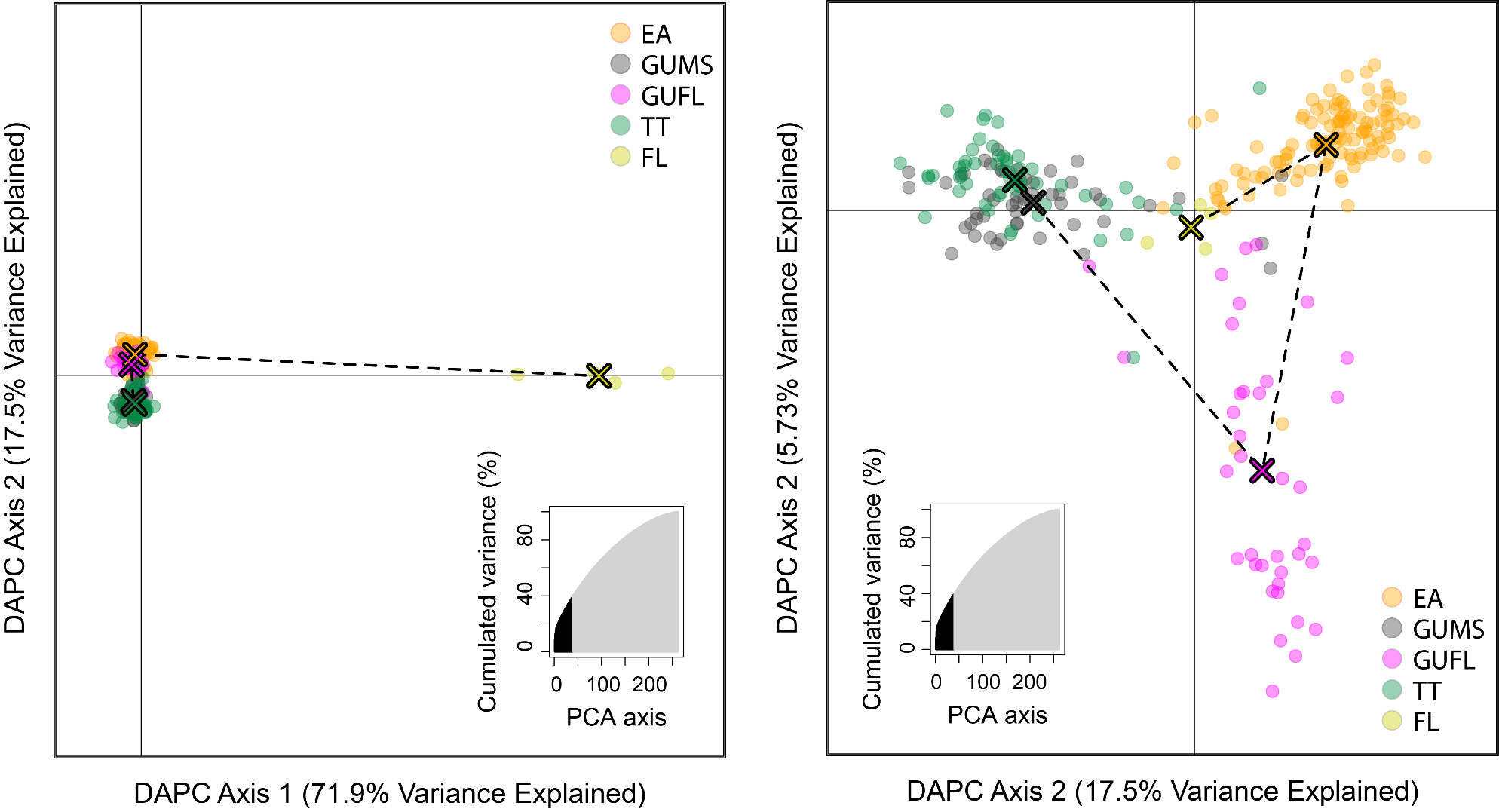

**Figure S5**: Discriminant Analysis of Principle Components (DAPC) for southeastern *Terrapene*. Each circle represents one individual, and each “X” delineates cluster centroids. The clusters (*K*=5, determined via Bayesian Information Criterion) represent: *T. c. carolina* (EA=Eastern), *T. c. major* (GU=Gulf Coast) from the Mississippi (GUMS) and Florida (GUFL) Panhandles, *T. m. triunguis* (TT=Three-toed), and *T. c. bauri* (FL=Florida). Inset plots demonstrate the number of retained principle components (PCs; N=40; shaded area), as determined using cross-validation (100 replicates with 90% of the dataset partitioned for training), versus the non-retained PCs (light gray area).

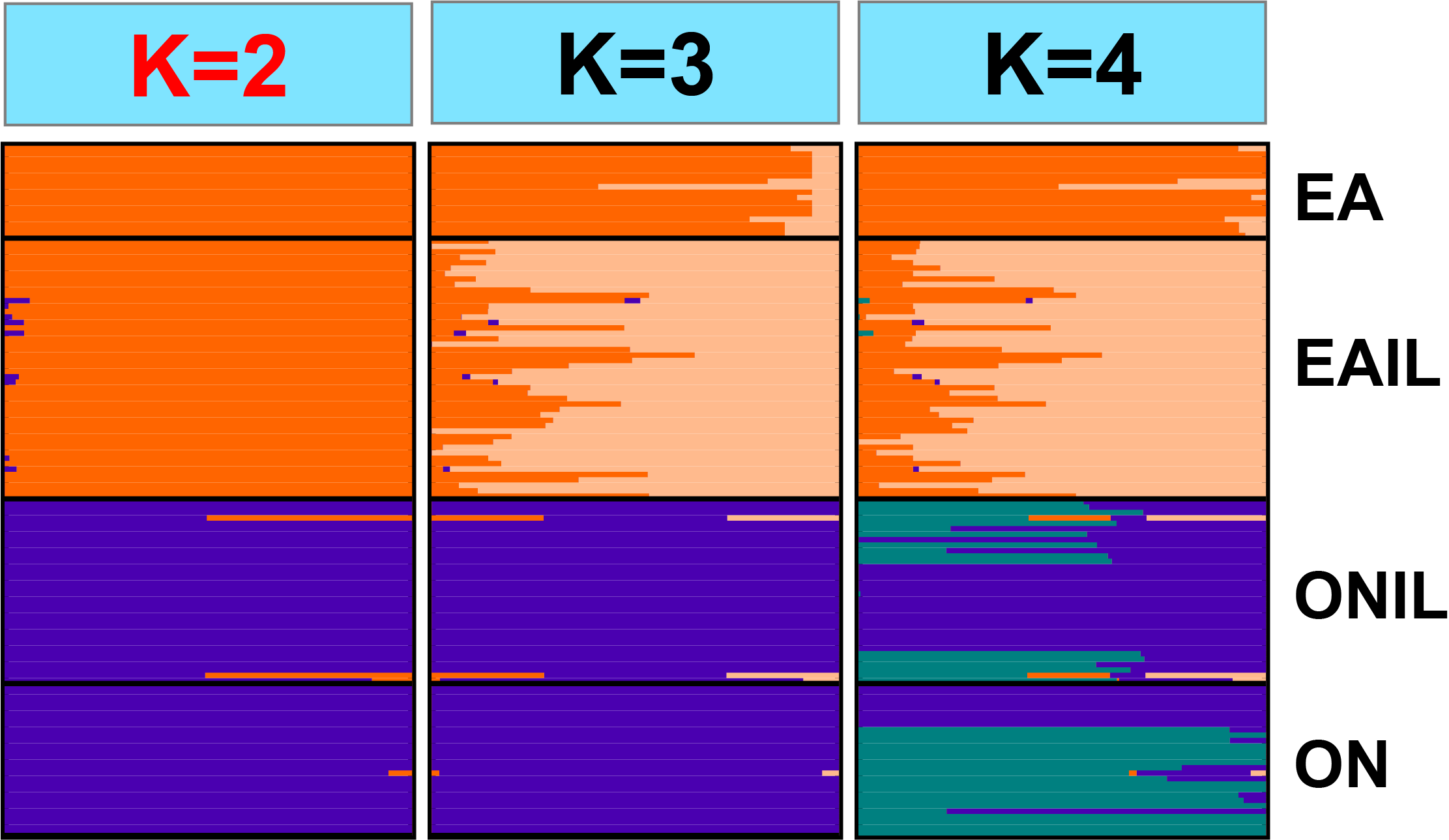

**Figure S6:** Top three midwestern *Terrapene* Admixture plots representing 10,338 unlinked ddRAD SNPs. The lowest cross-validation score was for *K*=2 (depicted in red at right), followed by *K*=4 then *K*=3. The first two letters of the population codes correspond to subspecific field identification (EA=Woodland, *T. c. carolina*; ON=Ornate, *T. o. ornata*). The second two letters (if present) represent locality codes for U.S. state (IL=Illinois). Populations lacking state locality code consist of multiple localities sampled outside the hybrid zone.

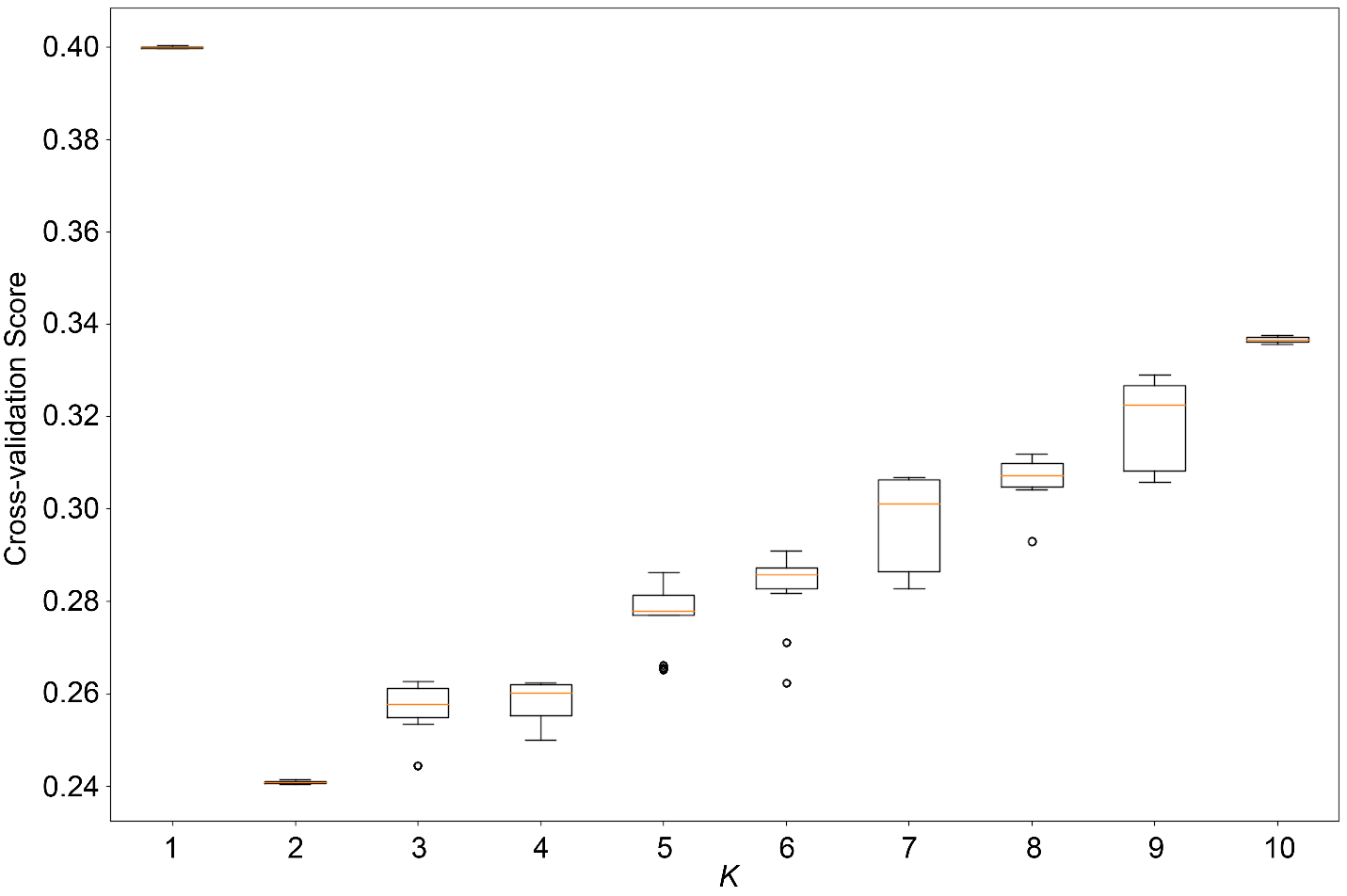

**Figure S7**: Cross-validation (CV) scores across all *K-values* (*K*=1-10) for Admixture runs containing samples from midwestern North America (N=135). Lower CV values indicate less error and stronger support for the corresponding *K*.

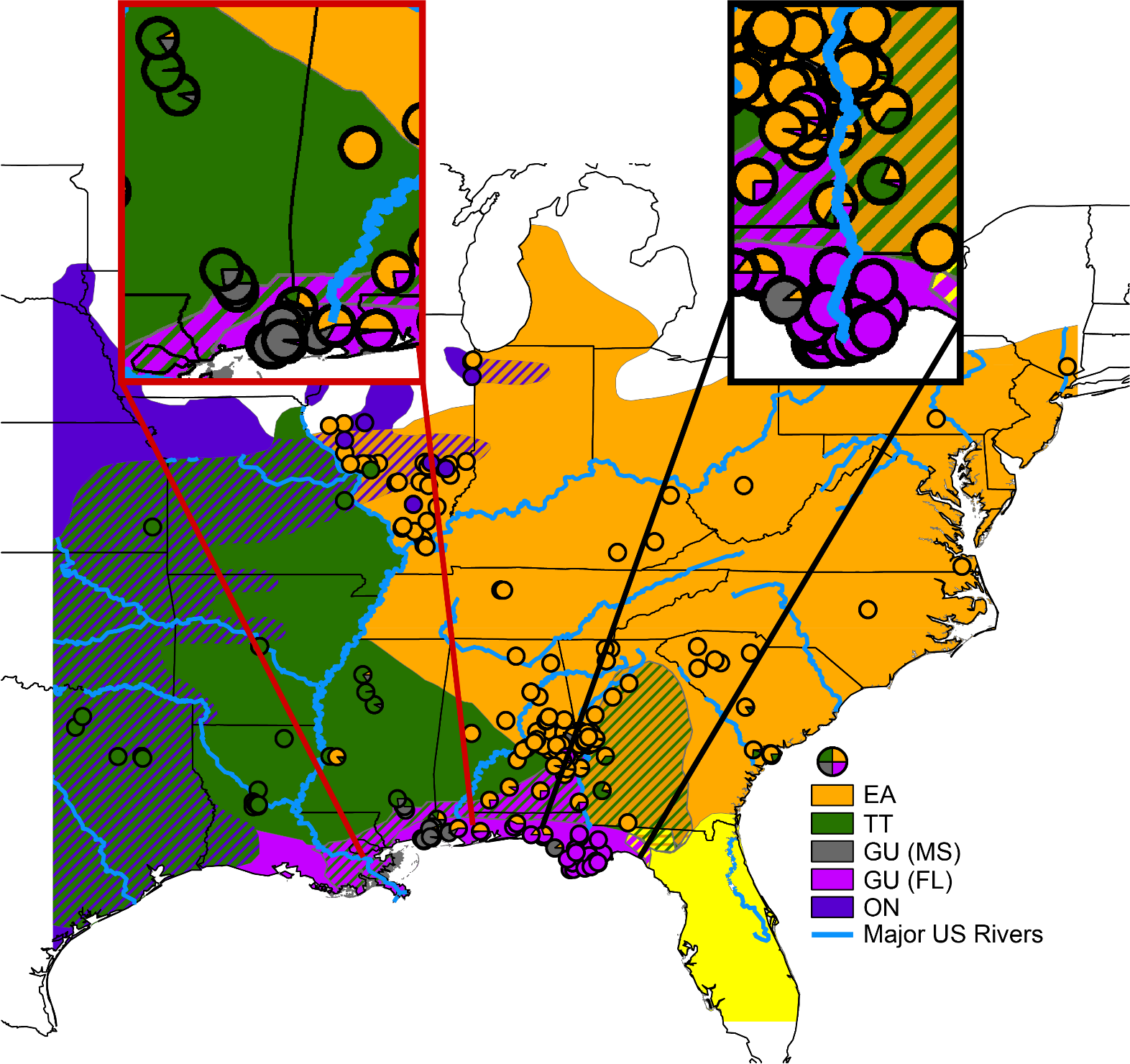

**Figure S8:** *Terrapene* distribution map. Cross-hatched areas represent contact zones. Circles indicate individual sampling localities, and the accompanying pie charts depict admixture proportions from the all-taxon *K*=5 (for midwestern individuals) and southeastern *K*=4 Admixture analyses (Fig. 1, 2). The expanded regions highlight two distinct *T. c. major* populations in the panhandles of Mississippi (red box) and Florida (black box), located in the Alabama and Apalachicola river basins, respectively. EA=Woodland (*T. carolina carolina*), GU=Gulf Coast (*T. c. major*), TT=Three-toed (*T. mexicana triunguis*), , ON=Ornate (*T. ornata ornata*).

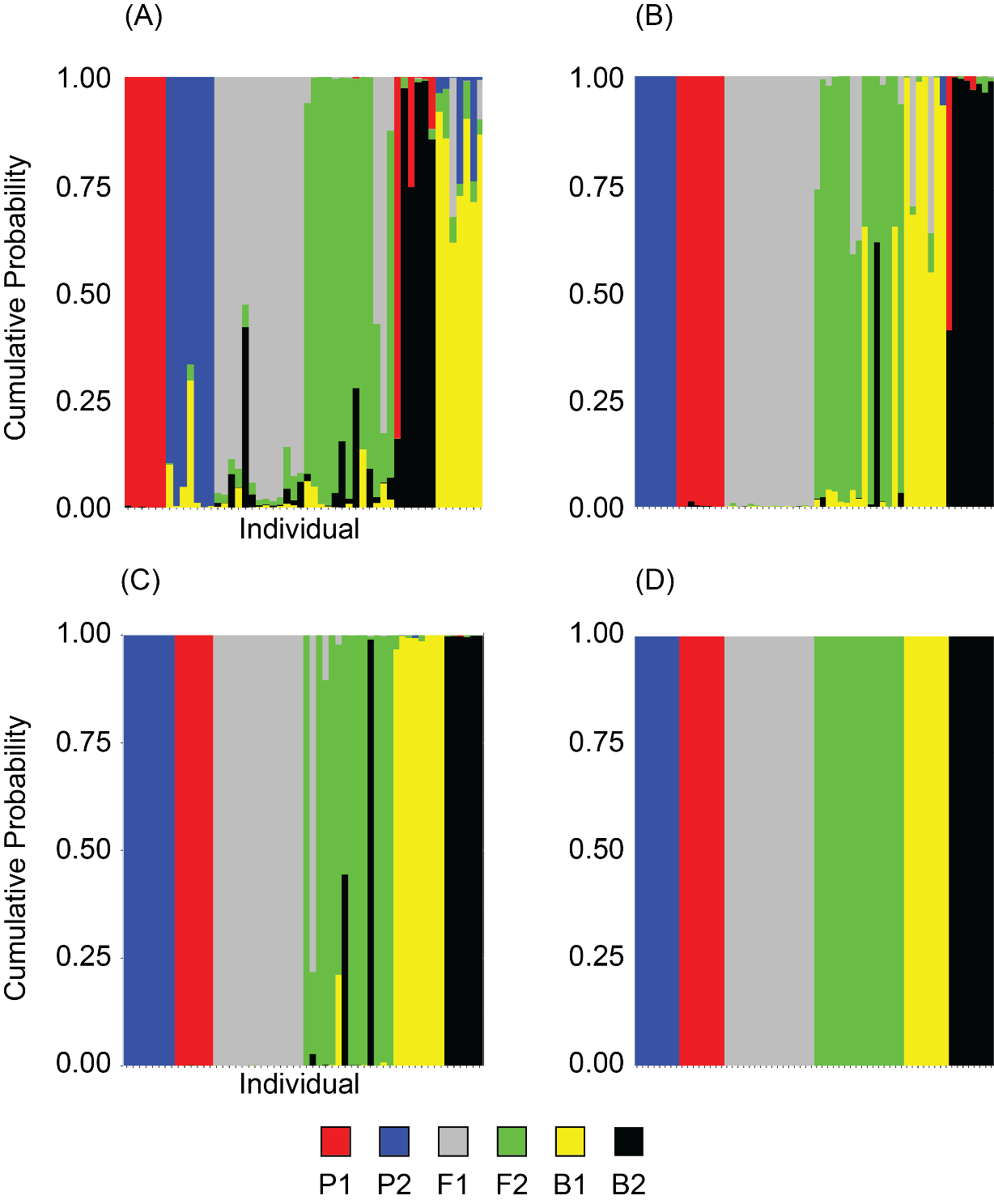

**Figure S9**: Results for *Terrapene* NewHybrids simulations that tested for convergence between inter- and intra-simulation replicates. Convergence was confirmed by the HybridDetective pipeline, thus only one of the virtually identical simulation replicates is presented for each of (A) *T. carolina carolina* (Woodland) X *T. c. major* (Gulf Coast), (B) *T. c. carolina* X *T. mexicana triunguis* (Three-toed), (C) *T. c. major* X *T. m. triunguis*, and (D) *T. c. carolina* X *T. o. ornata* (Ornate) is shown here. The genotype frequencies included parental groups (P_1_ and P_2_), first and second-generation hybrids (F_1_ and F_2_), and backcross (B_1_ and B_2_) generations.

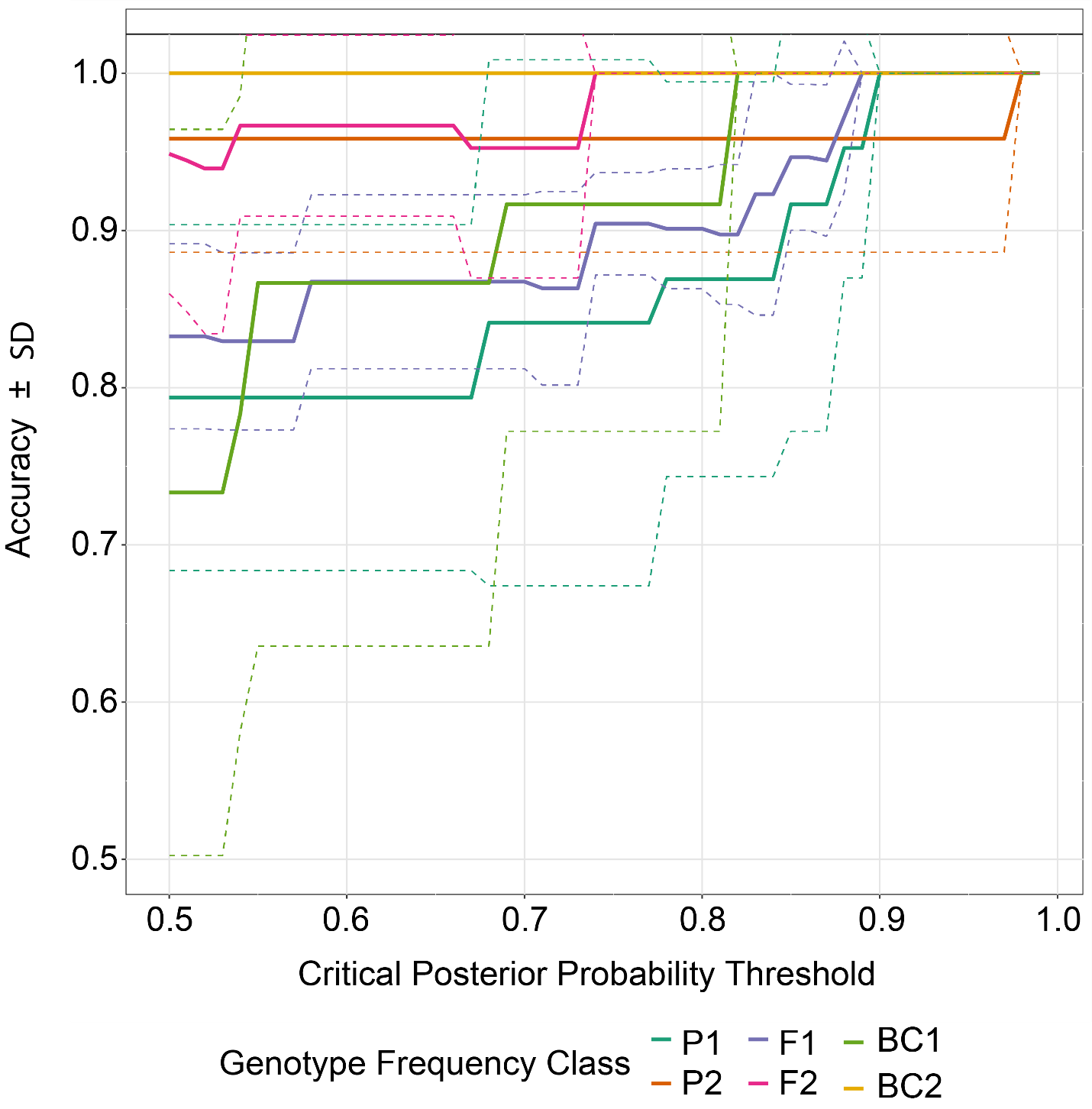

**Figure S10**: NewHybrids power analysis for *Terrapene carolina carolina* (Woodland) X *T. c. major* (Gulf Coast; EAxGU) showing predicted accuracy plotted against posterior probability thresholds. Accuracy was calcuated using simulated datasets for parental (P_1_ and P_2_), first and second-generation hybrid (F_1_ and F_2_), and backcross (B_1_ and B_2_) generations.

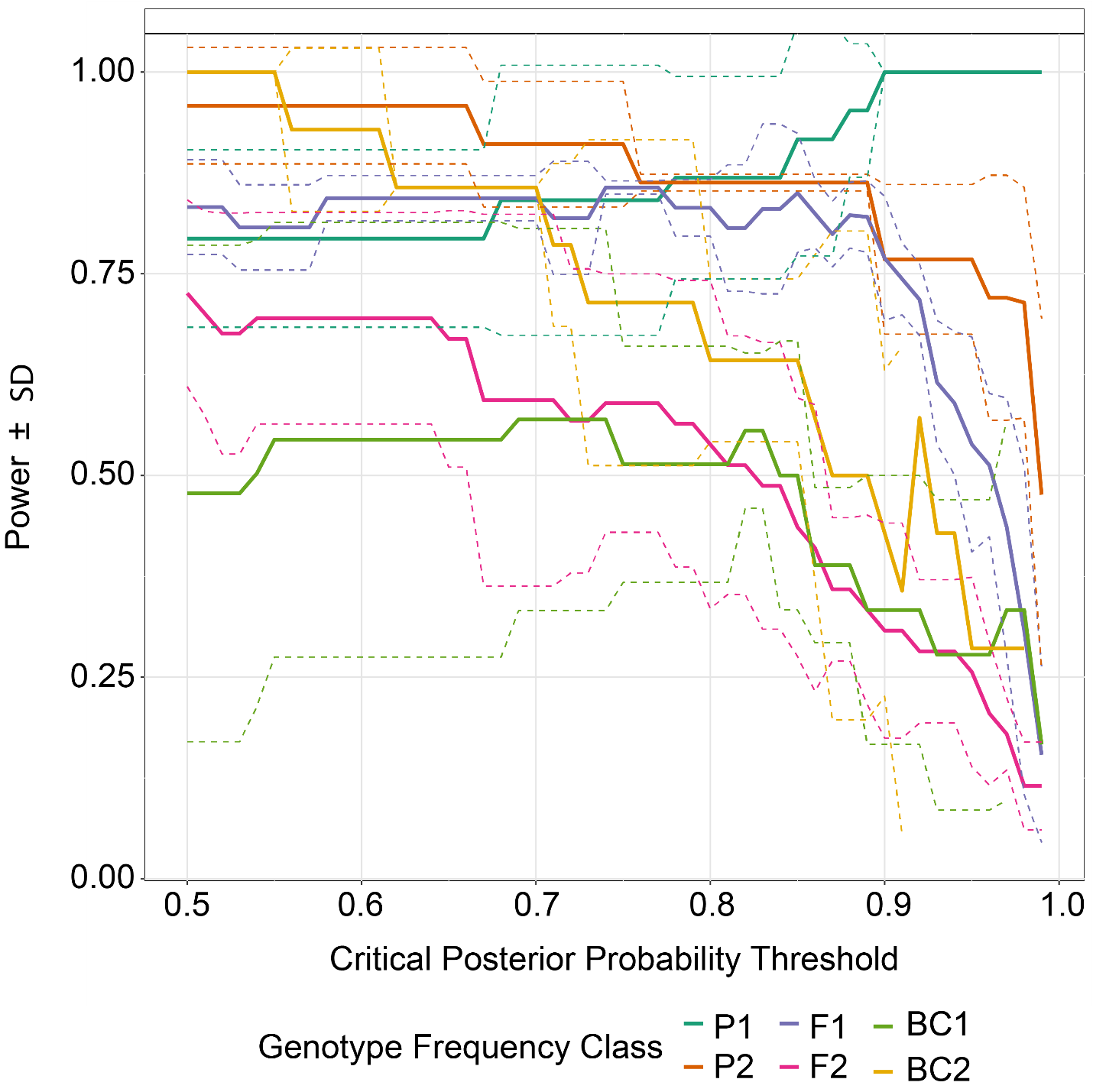

**Figure S11**: NewHybrids power analysis for *Terrapene carolina carolina* (Woodland) X *T. c. major* (Gulf Coast; EAxGU) showing predicted power plotted against posterior probability thresholds. Power was calcuated using simulated datasets for parental (P_1_ and P_2_), first and second-generation hybrid (F_1_ and F_2_), and backcross (B_1_ and B_2_) generations.

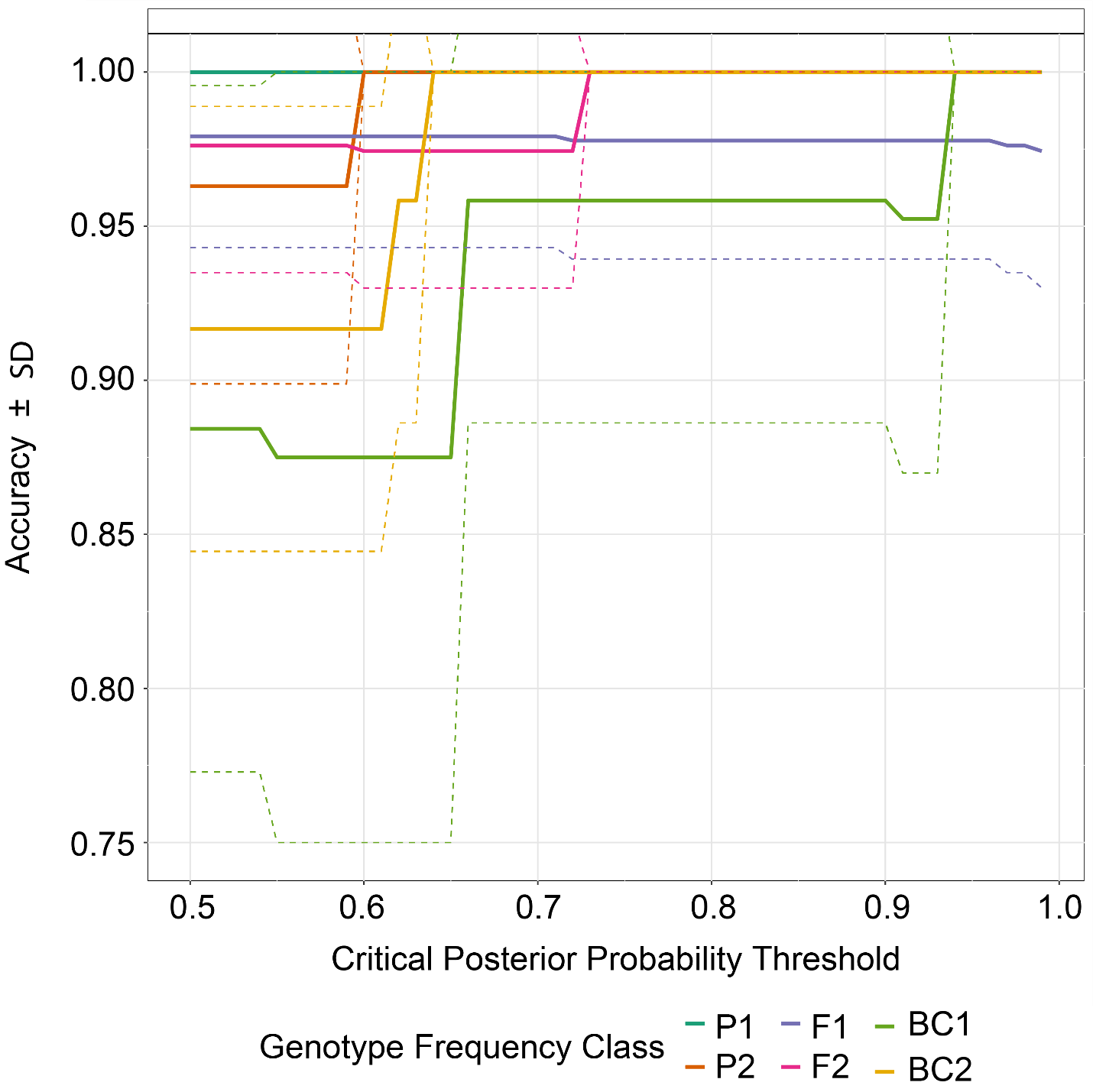

**Figure S12**: NewHybrids power analysis for *Terrapene carolina carolina* (Woodland) X *T. mexicana triunguis* (Three-toed; EAxTT) showing predicted accuracy plotted against posterior probability thresholds. Accuracy was calcuated using simulated datasets for parental (P_1_ and P_2_), first and second-generation hybrid (F_1_ and F_2_), and backcross (B_1_ and B_2_) generations.

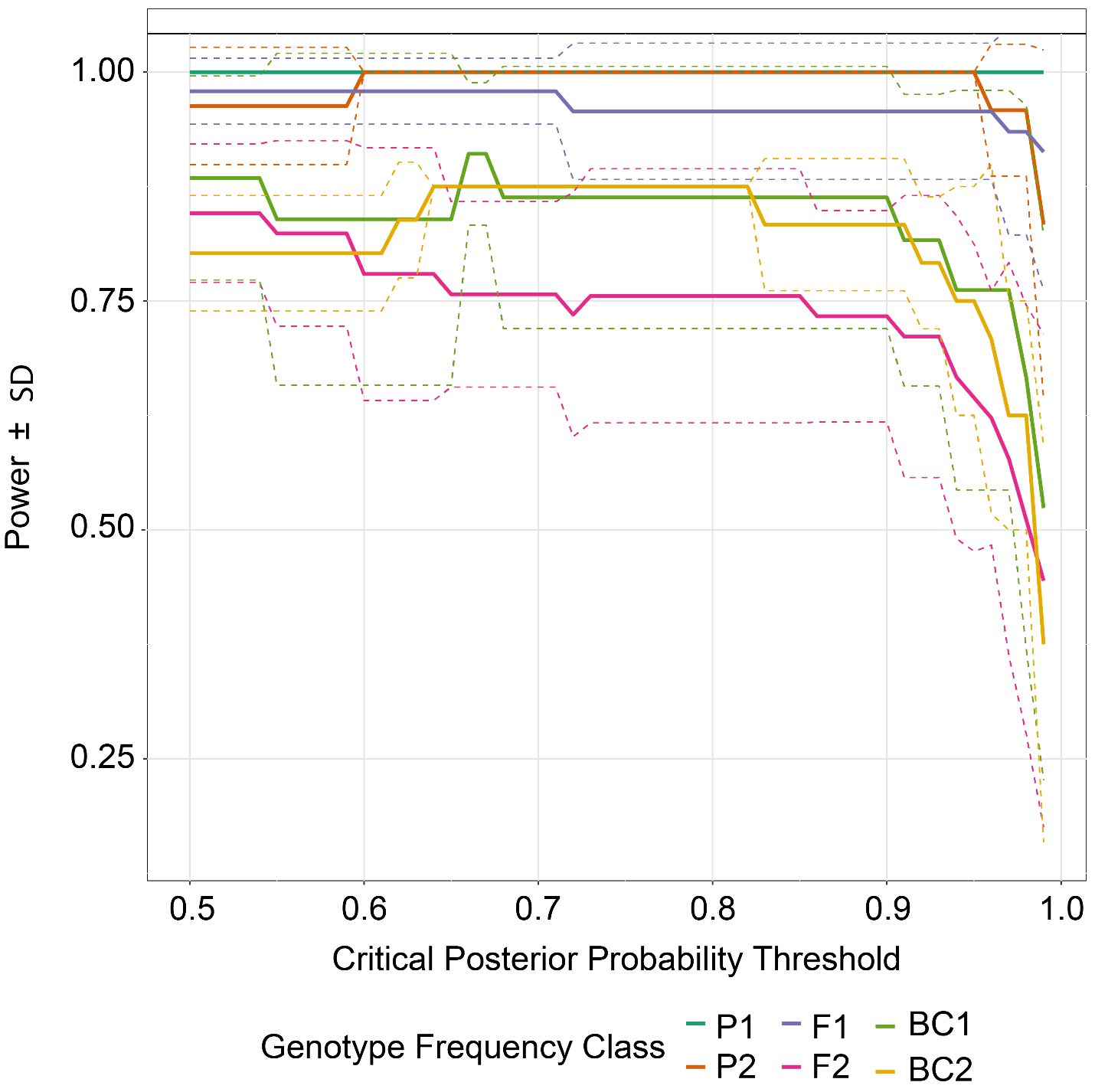

**Figure S13**: NewHybrids power analysis for *Terrapene carolina carolina* (Woodland) X *T. mexciana triunguis* (Three-toed; EAxTT) showing predicted power plotted against posterior probability thresholds. Power was calcuated using simulated datasets for parental (P_1_ and P_2_), first and second-generation hybrid (F_1_ and F_2_), and backcross (B_1_ and B_2_) generations.

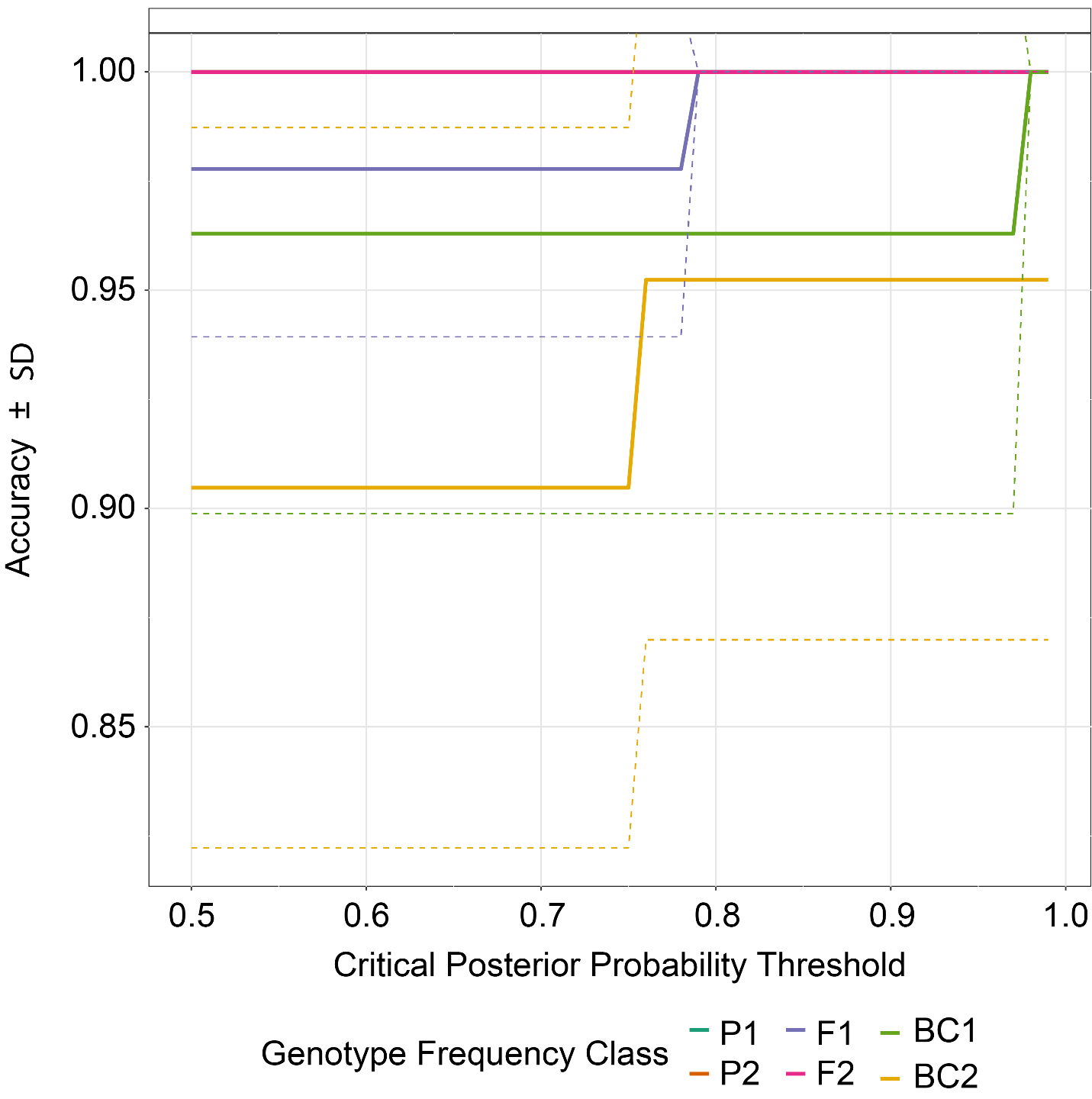

**Figure S14**: NewHybrids power analysis for *Terrapene carolina major* (Gulf Coast) X *T. mexicana triunguis* (Three-toed; GUxTT) showing predicted accuracy plotted against posterior probability thresholds. Accuracy was calcuated using simulated datasets for parental (P_1_ and P_2_), first and second-generation hybrid (F_1_ and F_2_), and backcross (B_1_ and B_2_) generations.

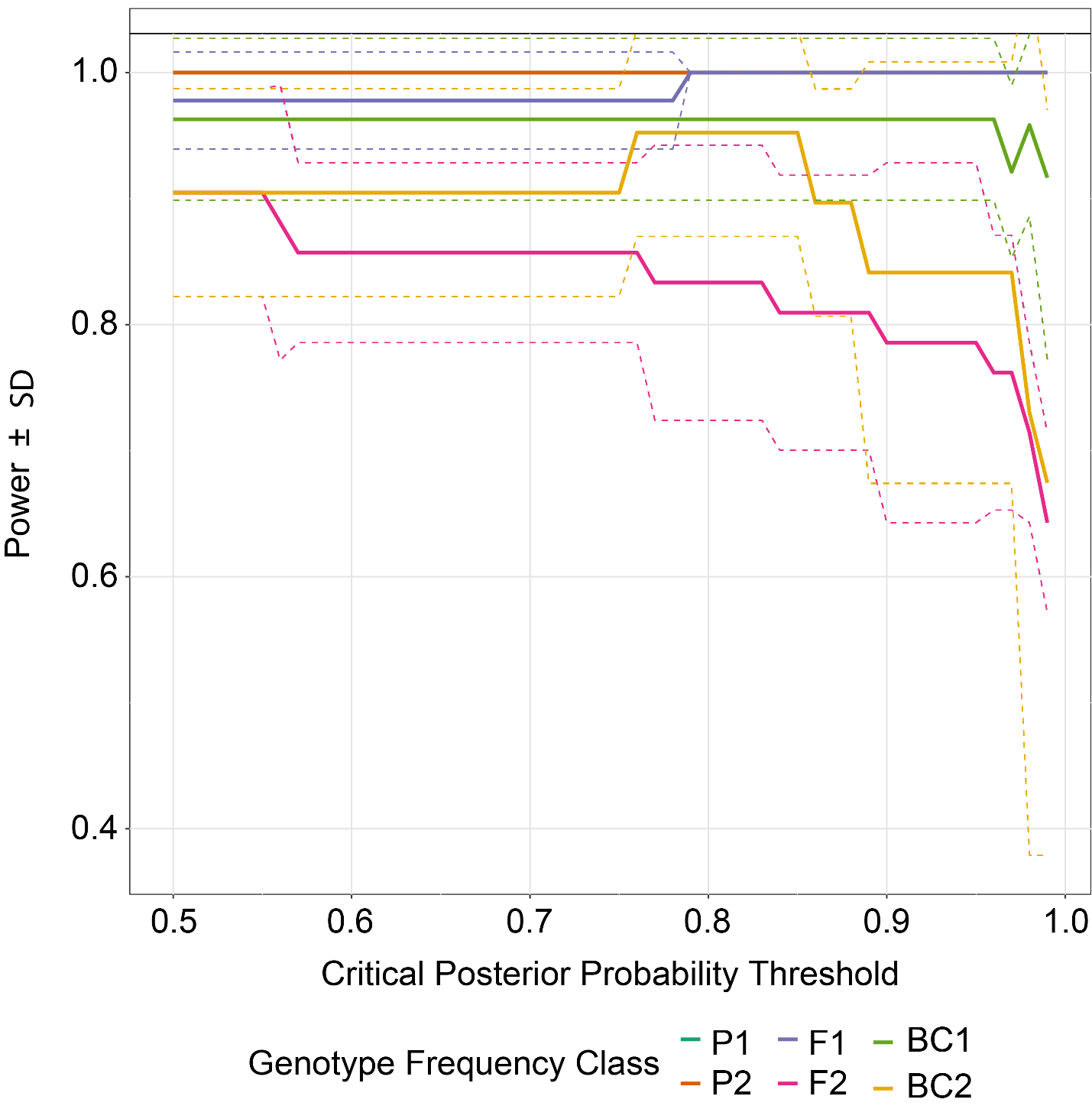

**Figure S15**: NewHybrids power analysis for *Terrapene carolina major* (Gulf Coast) X *T. mexicana triunguis* (Three-toed; GUxTT) showing predicted power plotted against posterior probability thresholds. Power was calcuated using simulated datasets for parental (P_1_ and P_2_), first and second-generation hybrid (F_1_ and F_2_), and backcross (B_1_ and B_2_) generations.

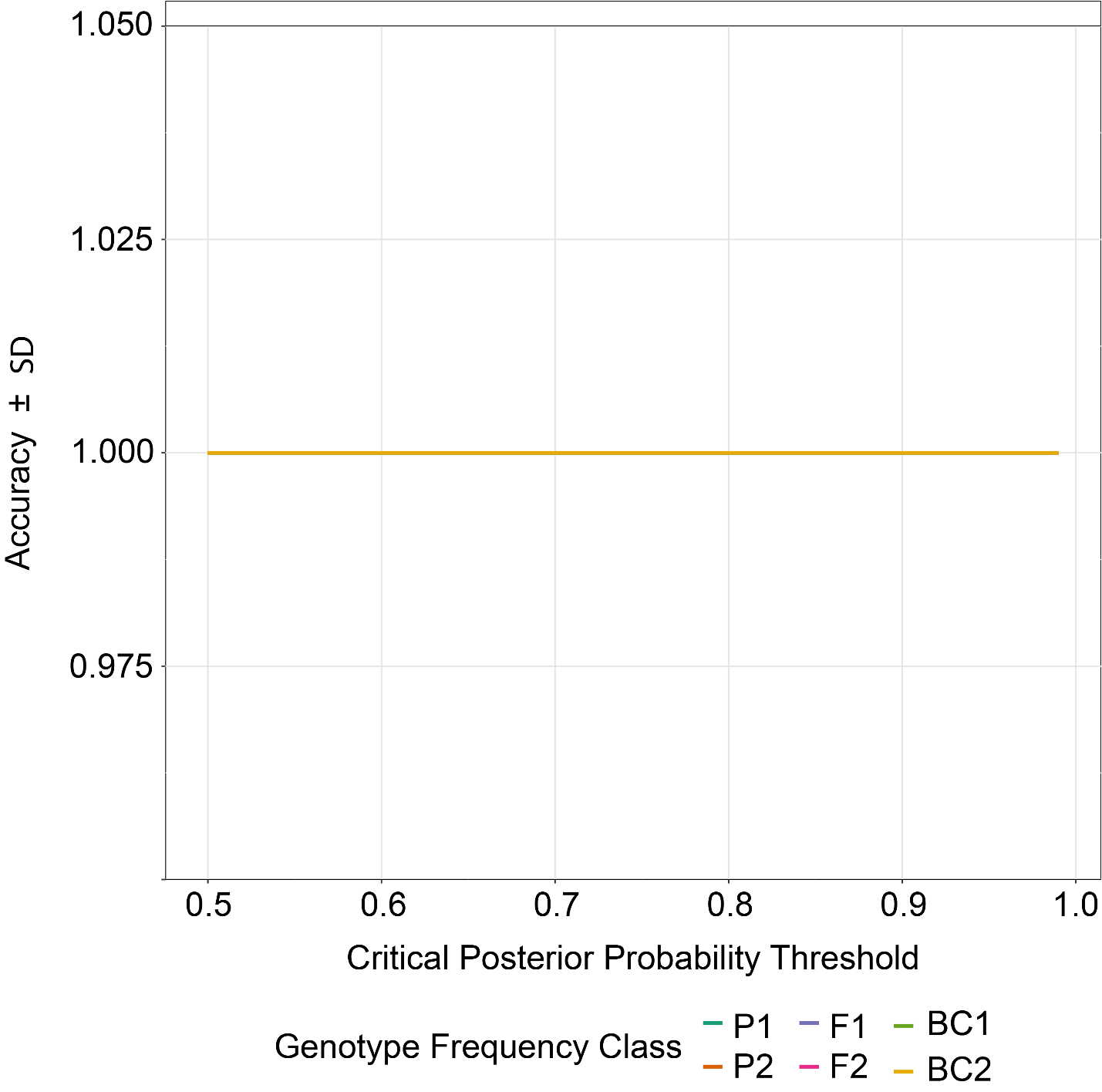

**Figure S16**: NewHybrids power analysis for *Terrapene carolina carolina* (Woodland) X *T. ornata ornata* (Ornate; EAxON) showing predicted accuracy plotted against posterior probability thresholds. Accuracy was calcuated using simulated datasets for parental (P_1_ and P_2_), first and second-generation hybrid (F_1_ and F_2_), and backcross (B_1_ and B_2_) generations.

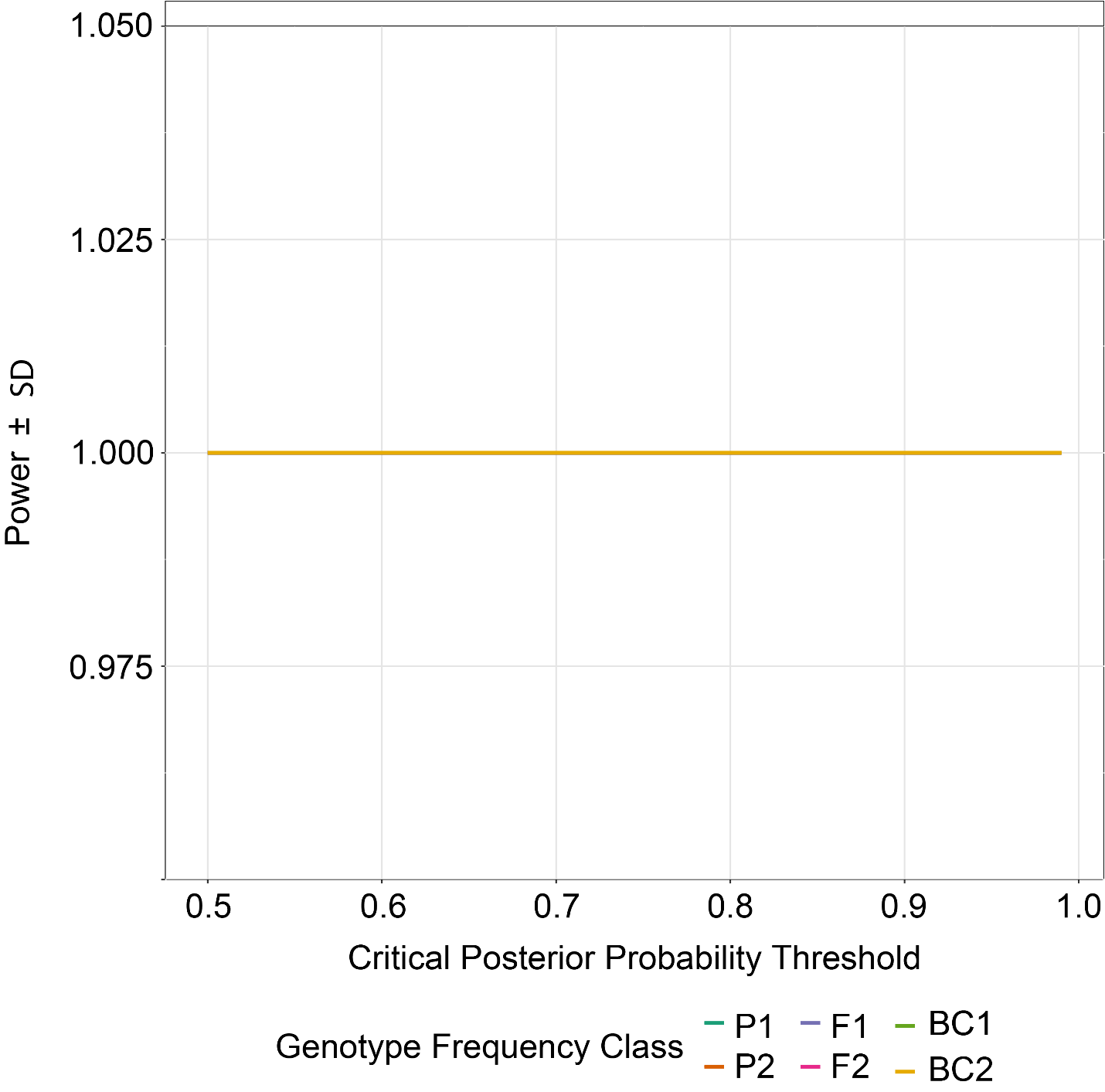

**Figure S17**: NewHybrids power analysis for *Terrapene carolina carolina* (Woodland) X *T. ornata ornata* (Ornate; EAxON) showing predicted power plotted against posterior probability thresholds. Power was calcuated using simulated datasets for parental (P_1_ and P_2_), first and second-generation hybrid (F_1_ and F_2_), and backcross (B_1_ and B_2_) generations.

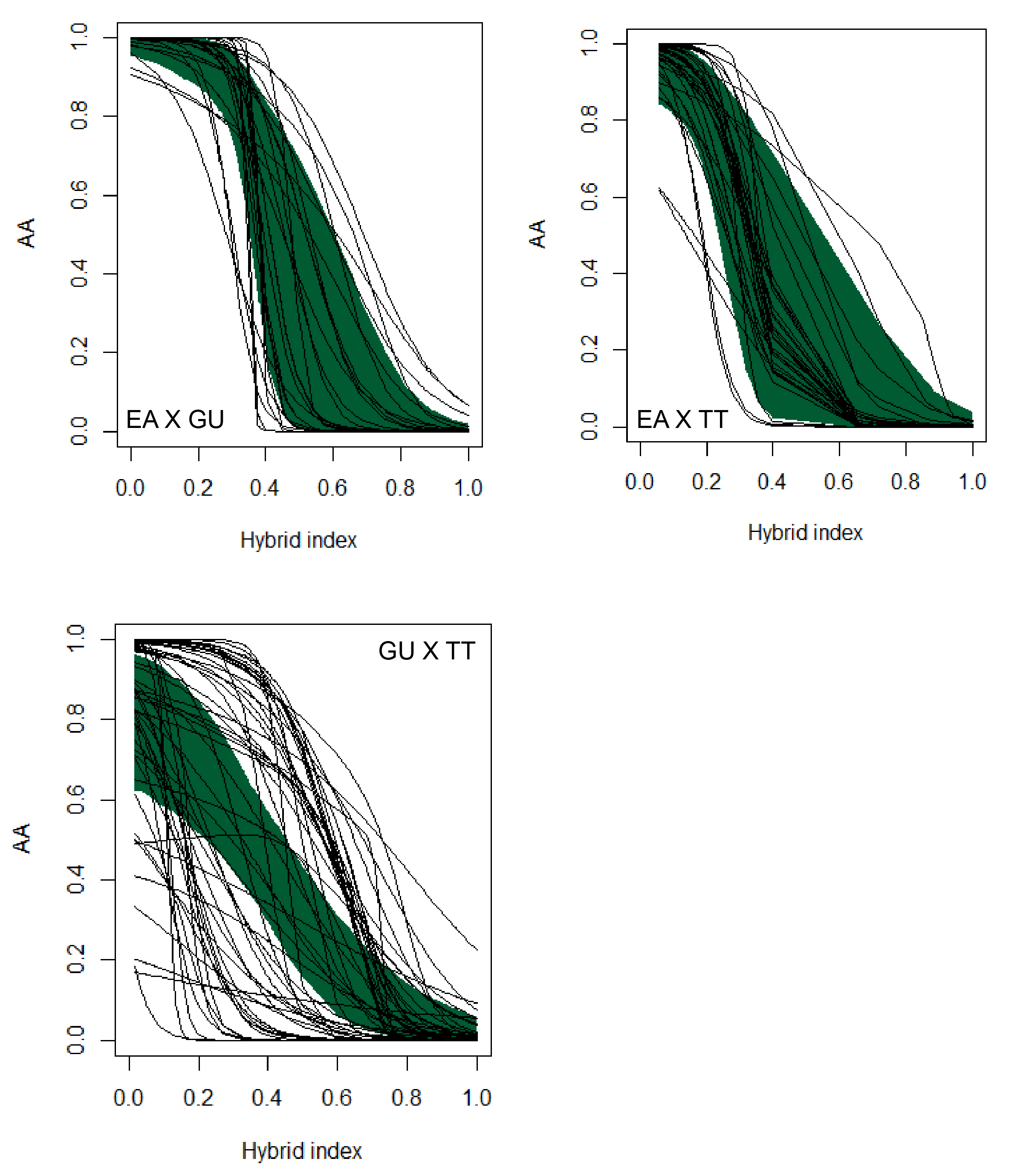

**Figure S18**: Genomic clines depicting outlier SNPs for all *Terrapene* ddRAD loci. Pairwise comparisons are between *T. carolina carolina* (EA=Woodland), *T. c. major* (GU=Gulf Coast), and *T. mexicana triunguis* (TT=Three-toed), with the number of loci per comparison being: N=10,106 (EAxGU); N=11,390 (EAxTT); and N=10,786 (GUxTT). The dark green area represents null expectations and each line is a genomic cline at one outlier locus. In each analysis, the P_1_ genotype represents EA, EA, and GU, respectively.
